# Molecular underpinning of intracellular pH regulation on TMEM16F

**DOI:** 10.1101/2020.06.15.153445

**Authors:** Pengfei Liang, Huanghe Yang

## Abstract

TMEM16F, a dual functional phospholipid scramblase and ion channel, is important in blood coagulation, skeleton development, HIV infection and cell fusion. Despite the advances in understanding its structure and activation mechanism, how TMEM16F is regulated by intracellular factors remains largely elusive. Here we report that TMEM16F lipid scrambling and ion channel activities are strongly influenced by intracellular pH (pHi). We find that low pH_i_ attenuates whereas high pH_i_ potentiates TMEM16F activation. Our biophysical characterizations pinpoint that the pH_i_ regulatory effects on TMEM16F stem from protonation and deprotonation of the Ca^2+^ binding sites, which in turn reduces and enhances Ca^2+^ binding affinity, respectively. We further demonstrate that intracellular alkalization of 0.5 pH_i_ can significantly promote TMEM16F activities in a choriocarcinoma cell line. Our findings thus uncover a regulatory mechanism of TMEM16F by pH_i_ and shine light on understanding the pathophysiological roles of TMEM16F in diseases with dysregulated pH_i_ including cancer.

## Introduction

The mammalian TMEM16 family consists of ten members. TMEM16A and TMEM16B are Ca^2+^-activated Cl^-^ channels (CaCCs), which participate in fluid secretion, smooth muscle contraction, gut motility, nociception, motor learning, anxiety and cancer (Hartzell, Putzier and Arreola, 2005; Caputo *et al*., 2008; Yang *et al*., 2008; Schroeder *et al*., 2008; Berg, Yang and Jan, 2012; Cho *et al*., 2012; Huang *et al*., 2012; Pedemonte and Galietta, 2014; Oh and Jung, 2016; Whitlock and Hartzell, 2017; Zhang *et al*., 2017; Crottès and Jan, 2019; Li *et al*., 2019). Majority of the other TMEM16 members are likely not CaCCs (Suzuki *et al*., 2010; Yang *et al*., 2012; Huang *et al*., 2013; Suzuki, Imanishi and Nagata, 2014; Whitlock *et al*., 2018; Bushell, Ashley C.W. Pike, *et al*., 2019). As one of the most studied TMEM16 proteins, TMEM16F is a dual functional Ca^2+^-activated non-selective ion channel and Ca^2+^-activated phospholipid scramblase (CaPLSase), which mediates phospholipids flip-flop across membrane lipid bilayer and rapidly destroy the asymmetric distribution of phospholipids on cell membranes (Suzuki *et al*., 2010; Yang *et al*., 2012). TMEM16F-mediated cell surface exposure of phosphatidylserine (PS), an amino-phospholipid concentrated in the inner leaflet of the plasma membrane, is essential for a number of cellular and physiological processes including blood coagulation (Suzuki *et al*., 2010; Yang *et al*., 2012), skeleton development (Ehlen *et al*., 2013; Ousingsawat *et al*., 2015), viral infection (Zaitseva *et al*., 2017), membrane microparticle release (Fujii *et al*., 2015), cell-cell fusion and placental development (Zhang *et al*., 2020). The loss-of-function mutations of human *TMEM16F* cause Scott Syndrome, a mild bleeding disorder characterized by a deficiency in CaPLSase-mediated PS exposure and subsequent defects on prothrombinase assembly, thrombin generation and blood coagulation (Suzuki *et al*., 2010; Castoldi *et al*., 2011). On the other hand, the TMEM16F deficient mice resist thrombotic challenges, suggesting that TMEM16F CaPLSase has the potential to serve as a promising therapeutic target for thrombotic disorders such as stroke, deep vein thrombosis and heart attack (Yang *et al*., 2012). Given its importance in health and disease, it is thus urgent to understand the molecular mechanisms and cellular functions of TMEM16F.

Recent structural and functional studies have advanced our understanding of the molecular architecture and the activation mechanism of TMEM16 CaPLSases (Pedemonte and Galietta, 2014; Brunner, Schenck and Dutzler, 2016; Whitlock and Hartzell, 2017; Falzone *et al*., 2018). The pore-gate domain of TMEM16 proteins consists of not only the permeation pathway for phospholipids and ions, but also two highly conserved Ca^2+^ binding sites (Yu *et al*., 2012; Brunner *et al*., 2014; Tien *et al*., 2014). Binding of intracellular Ca^2+^ (Ca^2+^_i_) triggers conformational changes, which lead to the opening of the activation gates and subsequent lipid and ion permeation (Dang *et al*., 2017; Paulino *et al*., 2017; Alvadia *et al*., 2019; Bushell, Ashley C. W. Pike, *et al*., 2019; Feng *et al*., 2019; T. Le *et al*., 2019). In addition to Ca^2+^_i_, membrane depolarization also facilitates the activation of TMEM16 CaCCs and TMEM16F ion channels (Yang *et al*., 2008, 2012; Dang *et al*., 2017; Paulino *et al*., 2017). In contrast to the comprehensive understanding of their activation mechanisms, the regulatory mechanisms of TMEM16 proteins just began to emerge.

Phosphatidylinositol (4,5)-bisphosphate (PIP_2_) was recently shown to plays a critical role in regulating TMEM16A and TMEM16F ion channel rundown or desensitization (Ta *et al*., 2017; De Jesús-Pérez *et al*., 2018; Ye *et al*., 2018; S. C. Le *et al*., 2019; Tembo *et al*., 2019; Yu *et al*., 2019). In addition, both intracellular and extracellular pH have also been reported to regulate endogenous CaCCs (Arreola, Melvin and Begenisich, 1995; Park and Brown, 1995; Qu and Hartzell, 2000) and heterologous expressed TMEM16A CaCCs (Chun *et al*., 2015; Cruz-Rangel *et al*., 2017; Segura-Covarrubias *et al*., 2020). The Oh laboratory recently reported that intracellular proton can inhibit TMEM16A CaCC by competing with Ca^2+^ on binding to the Ca^2+^ binding sites instead of affecting intracellular histidine residues (Chun *et al*., 2015). Nevertheless, it is not known if TMEM16F’s phospholipid scrambling and ion channel activities can also be regulated by pH_i_. Interestingly, TMEM16F is highly expressed in various tumor cells including the pancreatic ductal adenocarcinoma cells (Wang *et al*., 2018), glioma cells (Xuan, Wang and Xie, 2019), and choriocarcinoma BeWo cells (Zhang *et al*., 2020). Consistent with its high expression level, TMEM16F has been implicated in tumor cell proliferation, migration and metastasis (Jacobsen *et al*., 2013; Wang *et al*., 2018; Xuan, Wang and Xie, 2019). Given the fact that pH dysregulation (intracellular alkalization to pH_i_ 7.3-7.6 and extracellular acidification to pH 6.8-7.0) is one of the hallmarks of cancer (Webb *et al*., 2011; White, Grillo-Hill and Barber, 2017), it is thus critical to understand whether pH_i_ can regulate TMEM16F ion channel and CaPLSase activities.

In this study, we utilized patch clamp and fluorescence imaging methods to systematically characterize the impacts of pH_i_ on TMEM16F activation. Our results show that intracellular acidification attenuates whereas intracellular alkalization potentiates both TMEM16F ion channel and CaPLSase activities. pH_i_ mainly affects TMEM16F Ca^2+^-dependent activation with negligible effect on its voltage-dependent activation. Our biophysical analysis and mutagenesis studies further demonstrates that the pH_i_ regulatory effects on TMEM16F stem from the protonation and deprotonation of the Ca^2+^ binding residues, which in turn reduces and enhances Ca^2+^ binding affinity, respectively. We also show that intracellular alkalization by 0.5 pH_i_ from physiological pH_i_ significantly promotes TMEM16F ion channel and CaPLSase activities in BeWo choriocarcinoma cells. Our findings thus uncover a new regulatory mechanism of TMEM16F, which will facilitate our understanding of the physiological and pathological functions of TMEM16F and other TMEM16 family members in health and disease.

## Results

### pH_i_ regulates both TMEM16A and TMEM16F ion channel activities

In order to evaluate pH_i_ effect on TMEM16F ion channel activity, we first used TMEM16A CaCC to establish an optimized electrophysiology protocol for quantification. Instead of using the gap-free protocol at fixed membrane voltages in a previous study (Chun *et al*., 2015), we used inside-out patches to measure TMEM16A current activation across a wide range of voltages under different pH_i_ so that the conductance-voltage (*G-V*) relationships can be constructed and the pH_i_ effects are compared at different pH_i_, voltages and Ca^2+^_i_. Consistent with the previous report (Chun *et al*., 2015), our analysis also shows that low pH_i_ (=6.1) greatly suppresses whereas high pH_i_ (=8.9) significantly potentiates TMEM16A activation in the presence of 0.5 µM Ca^2+^_i_ as evidenced by the dramatic leftward and rightward shifts of the *G-V* curves, respectively (Fig.1, A, B). By plotting the mean conductance under different pH_i_ and voltages, which are normalized to the maximum conductance under +100 mV and at pH_i_=8.9, we constructed the conductance-pH_i_ (*G-pH*_*i*_) relation (Fig. 1C). By fitting the *G-pH*_*i*_ relation with linear regression, we obtained the slopes of ∼0.3 for activation voltages from +20 to +100 mV. The slopes of the *G-pH*_*i*_ plots represent the apparent pH_i_ sensitivity under 0.5 µM Ca^2+^_i_, which indicates that each pH_i_ unit change or every 10-fold change of intracellular H^+^ concentration from physiological pH_i_ can lead to ∼30% change of TMEM16A channel activation across different activation voltages. The almost identical slopes of the *G-pH*_*i*_ plots under different membrane potentials suggest that pH_i_ has negligible effect on TMEM16A voltage-dependent activation (Fig. 1C).

**Figure 1.**
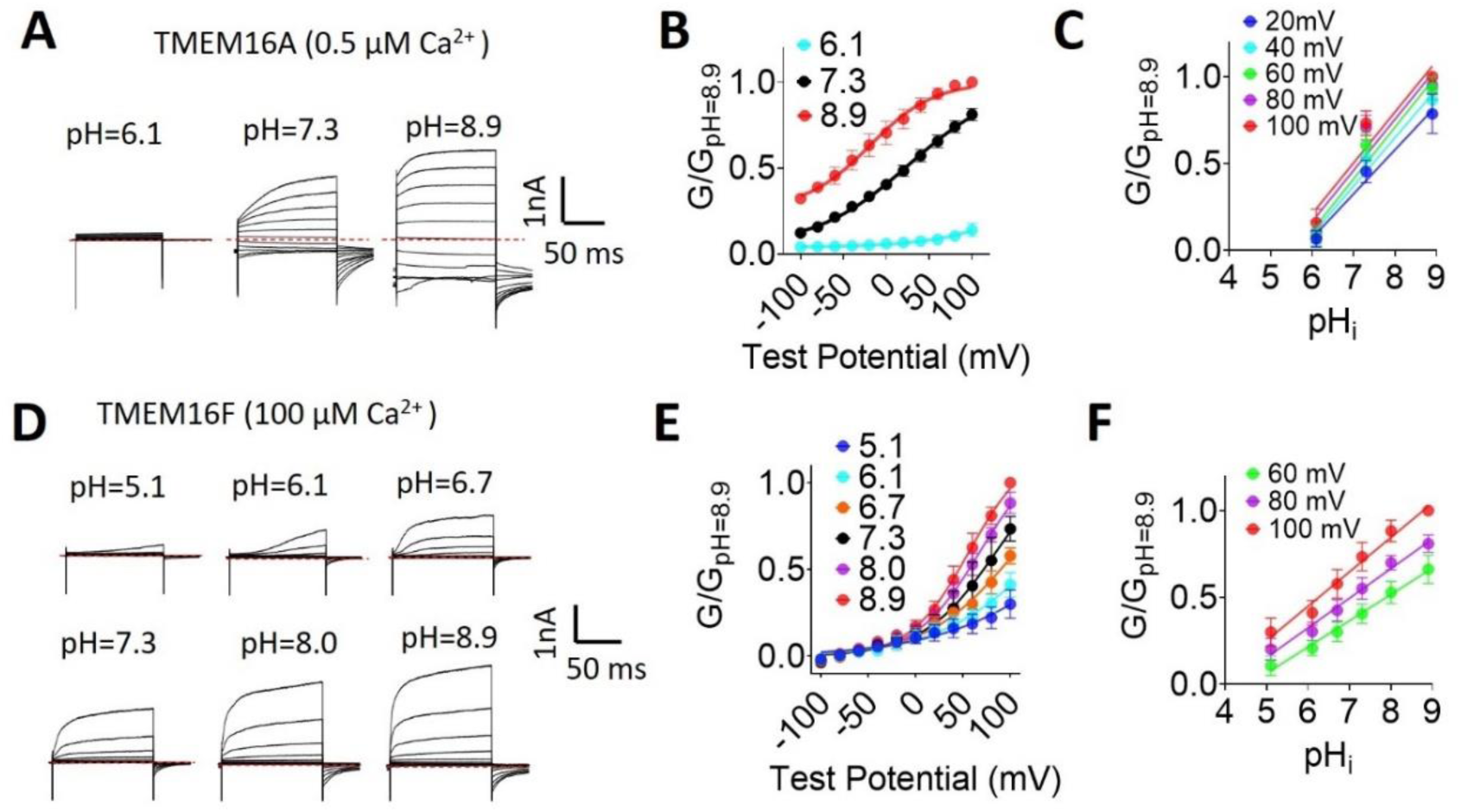
pH_i_ regulates both TMEM16A and TMEM16F ion channel activities. (**A)** Representative TMEM16A currents recorded from inside-out patches perfused with intracellular solutions containing 0.5 µM Ca^2+^ at different pH_i_. Currents were elicited by voltage steps from −100 to +100 mV with 20 mV increments. The holding potential was −60 mV. All the traces shown were from the same patch. (**B)** Mean *G-V* relations of the TMEM16A channels under different pH_i_ at 0.5 µM Ca^2+^. Relative conductance was determined by measuring the amplitude of tail currents 400 μs after repolarization to a fixed membrane potential (−60 mV). The smooth curves represent Boltzmann fits *G/G*_*max*_*=1/(1+exp(−ze(V−V*_*1/2*_*)/kT). G*_*max*_ is tail current amplitude in response to depolarization to +100 mV in 0.5 μM Ca^2+^ at pH_i_=8.9. Error bar represents SEM (n=7). **(C)** Mean conductance of TMEM16A at different pH_i_ was normalized to the maximum conductance at pH_i_=8.9 at different voltages and then plotted as a function of pH_i_ (*G-pH*_*i*_ relationship). Data were fitted with linear regression and the mean slopes from fittings were 0.27,0.28, 0.29,0.3 and 0.29 for 20, 40, 60, 80 and 100 mV, respectively (n=7). **(D)** Representative TMEM16F currents recorded from inside-out patches perfused with intracellular solutions containing 100 µM Ca^2+^ at different pH_i_. Currents were evoked by voltage steps from −100 to +100 mV with 20 mV increments and the holding potential was −60 mV. All traces shown were from the same patch. **(E)** Mean *G-V* relations of TMEM16F current under different pH_i_. The smooth curves represent Boltzmann fits similar as in **(B)**. Error bar represents SEM (n=7). (**F)** *G-pH*_*i*_ relationship for TMEM16F channels. Data were fitted with linear regression and the mean slopes from fittings were 0.19, 0.20 and 0.21 for 60, 80 and 100 mV, respectively (n=7).

Using the same patch clamp protocol, we next examined whether pH_i_ can regulate TMEM16F ion channel activity. As TMEM16F is less sensitive to Ca^2+^ than TMEM16A (Yang *et al*., 2012), we activated TMEM16F channels stably expressed in HEK293 cells using 100 µM Ca^2+^_i_ and membrane depolarization. Our inside-out patch clamp recordings show that pH_i_ also regulates TMEM16F channel activation as it does to TMEM16A (Fig. 1D, E). Alkalized pH_i_ (8.0 and 8.9) greatly potentiates TMEM16F current compared to a physiological pH_i_ of 7.3. In contrast, acidic pH_i_ (5.1, 6.1 and 6.7) suppresses TMEM16F channel activation in a pH_i_-dependent fashion. The apparent pH_i_ sensitivity of TMEM16F channel was assessed by plotting the *G-pH*_*i*_ relationship (Fig. 1F). Our linear regression fittings result in a slope range from 0.19 to 0.21, indicating that for each pH_i_ unit reduction, TMEM16F channel activation will be reduced by ∼20%. Similar to TMEM16A, TMEM16F pH_i_ sensitivity is also voltage independent as evidenced by the parallel *G-pH*_*i*_ relationships at different voltages. By extrapolating the fittings of the *G-pH*_*i*_ relationships, we predict that TMEM16F channel activity would be completely abolished when pH_i_ drops below 4. As TMEM16F channel subjects to PIP_2_-dependent rundown or desensitization under inside-out configuration (Ye *et al*., 2018), we thus estimated the influence of rundown on our measurements of the pH_i_ effects. The peak currents elicited by a voltage-step protocol only reduced ∼20% (Fig. S1A-B) within the time window to complete all the recordings shown in Fig.1D (about 1 minute). This result suggests that TMEM16F channel rundown only has small effects on our quantification of the pH_i_ effects. It is worth mentioning that we sequentially perfused different pH_i_ solutions from low pH_i_ to high pH_i_ and the normalized TMEM16F current to the maximum current in pH_i_ 8.9 (Fig. 1E). Thus, if taken channel rundown into account, the pH_i_ sensitivity of TMEM16F channel in our measurements would have been underestimated. Taking together, our patch clamp recordings reveal that pH_i_ is an important intracellular factor to regulate both TMEM16A and TMEM16F ion channel activities.

### pH_i_ regulates TMEM16F scrambling activity

Having shown that pH_i_ can regulate TMEM16F ion channel activity, we next sought to examine whether pH_i_ can also influence TMEM16F lipid scrambling activity. In order to quantify TMEM16F lipid scrambling in different pH_i_, we established a modified fluorescence imaging assay (Le, Le and Yang, 2019; T. Le *et al*., 2019) to overcome the difficulties in precisely controlling pH_i_ and Ca^2+^_i_ (Fig. 2A). In this lipid scrambling fluorometry assay, pH_i_ and Ca^2+^_i_ are accurately controlled by solution exchange between the glass pipettes and the cytosol of the whole-cell patch clamped cells. Once we break the patch membrane into whole-cell configuration, the pipette solution with fixed pH_i_ and Ca^2+^_i_ starts to diffuse into the cytosol and activate TMEM16F CaPLSases. Fluorescently conjugated Annexin V (AnV-CF 594) is simultaneously perfused to the patched cell. After a delay during which Ca^2+^_i_ reaches the threshold to activate TMEM16F CaPLSase, AnV starts to be attracted to cell surface by the externalized PS (Fig. 2B). The fluorescence signal accumulated on cell surface increases over time and exhibit a sigmoid relationship with time (Fig. 2C). By fitting the curve with a generalized logistic function, we can obtain t_1/2_, the time needed to reach half-maximum fluorescence when the macroscopic CaPLSase activity reaches maximum speed. At physiological pH_i_ of 7.3, 100 µM Ca^2+^_i_ activates TMEM16F lipid scramblases with an averaged t_1/2_ of 20.3±2.4 minutes (Fig. 2B-D and Movie S1). We found that intracellular alkalization (pH_i_=8.9) markedly shortened t_1/2_ to about 11.6±3.5 minutes. In stark contrast, intracellular acidification (pH_i_=6.1) significantly prolonged t_1/2_ to 29.1±3.0 minutes. The changes of t_1/2_ under different pH_i_ indicate that low pH_i_ attenuates and high pH_i_ enhances TMEM16F CaPLSase activities. We also quantified the maximum lipid scrambling rate at t_1/2_ or the slope (k) of the ‘linear phase’ on the sigmoid curves under different pH_i_. We found that low pH_i_ (pH_i_=6.1) significantly reduced slope k from 0.26±0.04 at physiological pH_i_ to 0.19±0.04, whereas high pH_i_ of 8.9 increased slope k to 0.4±0.02 (Fig. 2E), suggesting that pH_i_ affects the maximum lipid scrambling rate of TMEM16F. To ensure the enhanced PS externalization at high pH_i_ is mediated by TMEM16F CaPLSases instead of through other mechanisms such as alkalization induced apoptosis (Lagadic-Gossmann, Huc and Lecureur, 2004), we used the same assay to test our TMEM16F knockout (KO) HEK293 cells (Le, Le and Yang, 2019) in pH_i_ 8.9. In stark contrast to fast lipid scrambling in TMEM16F stable HEK293 cells (Fig. 2C-D), high pH_i_ did not induce AnV binding to the TMEM16F KO cells over time course of 12.5 minutes (Fig. S1C-D and Movie S2). This experiment suggests that apoptosis-induced PS exposure is unlikely to contribute to the pH_i_ effects on TMEM16F CaPLSases under our experimental conditions.

**Figure 2.**
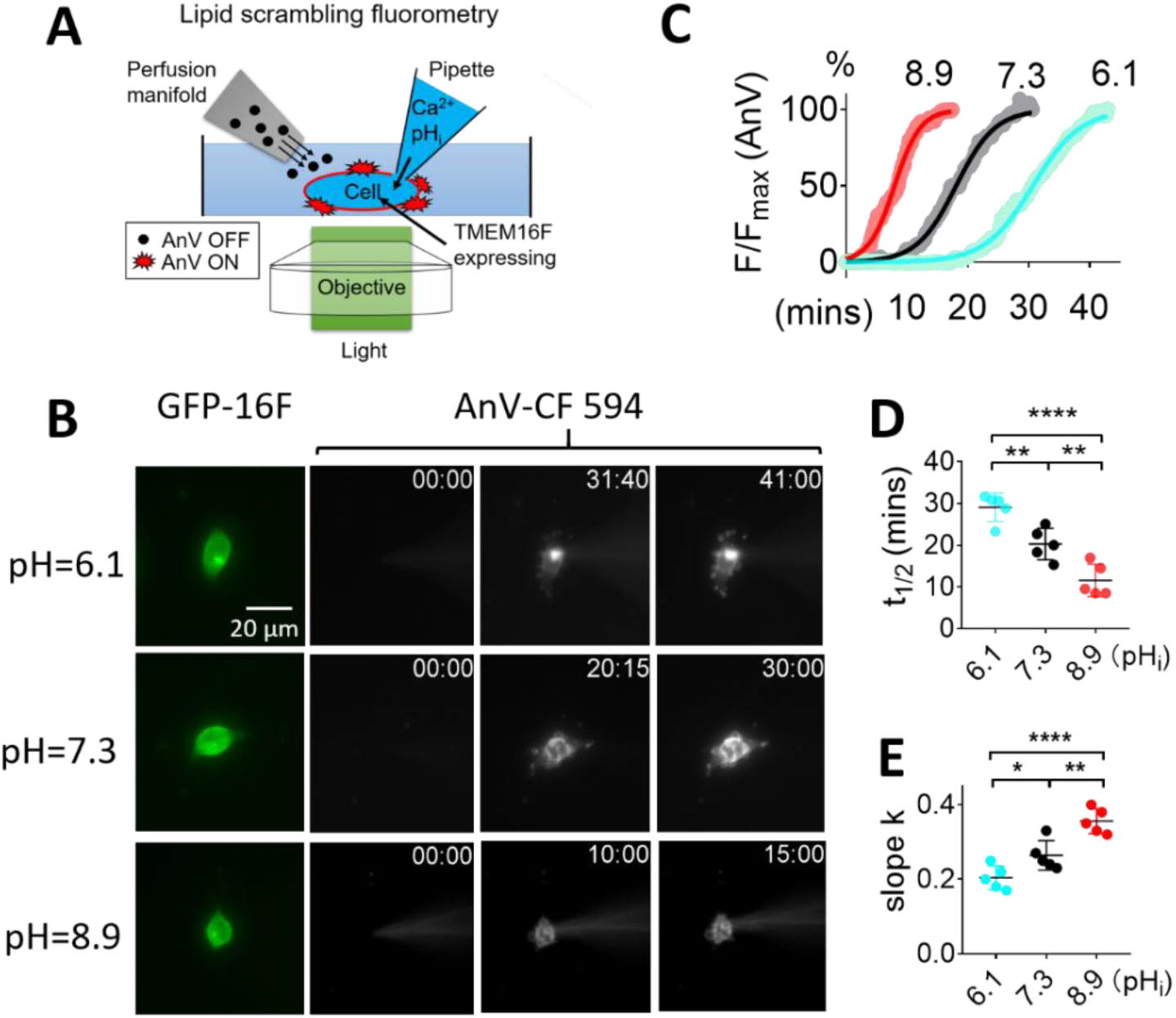
pH_i_ regulates TMEM16F lipid scrambling activity. **(A)** Schematic design of the lipid scrambling fluorometry assay. CaPLSase activity is monitored by cell surface accumulation of fluorescently tagged Annexin-V (AnV), a phosphatidylserine (PS) binding protein that is continuously perfused through a perfusion manifold. AnV fluorescence remains dim in bulk solution and will strongly fluoresces after being recruited by cell surface PS, which is externalized by CaPLSases. We utilize whole-cell patch pipettes to deliver intracellular solutions into the cytosol to achieve precise control of intracellular pH and Ca^2+^. Once breaking into whole-cell configuration, the pipette solution will rapidly diffuse into the cell and activate CaPLSases. The AnV fluorescence signal on cell surface will be continuously recorded with 5-second interval. **(B)** Representative lipid scrambling fluorometry images of the HEK293T cells stably expressed with TMEM16F-eGFP (left, green signal) at different pH_i_. For the AnV-CF 594 signal on the right, the first column is fluorescence signal immediately after forming whole-cell configuration; the second column is the time when fluorescence intensity reached half maximum (t_1/2_) and the last column is the time when fluorescence signal roughly reach plateau. The time point (minutes followed by seconds) of each image after breaking into whole-cell configuration are shown on the top right corner. The pipette solution contained 100 µM Ca^2+^ and holding potential was −60 mV. **(C)** The time course of AnV fluorescence signal at different pH_i_ as shown in (B). The AnV signal was normalized to the maximum AnV fluorescence intensity at the end of each recording. The smooth curves represent fits to generalized logistic equation, *F=F*_*max*_*/[1+exp(-k(t-t*_*1/2*_))]. **(D)** Under 100 µM Ca^2+^, t_1/2_ for pH_i_ = 6.1,7.3 and 8.9 are 29.1±3.0 minutes; 20.3±2.4 minutes and 11.6±3.5 minutes (n=5), respectively. **(E)** Under 100 µM Ca^2+^, slopes for pH_i_ = 6.1,7.3 and 8.9 are 0.19±0.04, 0.26±0.04 and 0.40±0.02 (n=5), respectively. Values represent as mean ± SEM. *=p<0.05, **=p<0.01 and **** = p < 0.0001, using one-way ANOVA followed by Tukey’s test.

Taken together, our results show that pH_i_ regulates both TMEM16F ion channel and scrambling activities. In comparison to physiological pH_i_, intracellular acidification suppresses TMEM16F activation, whereas intracellular alkalization enhances TMEM16F activation. This implies that the pH_i_ effects on TMEM16F ion channel and lipid scrambling might share the same molecular mechanism.

### pH_i_ has no effect on voltage-dependent activation of TMEM16A and TMEM16F channels

We next set out to dissect the molecular mechanism underlying pH_i_ regulation on TMEM16 ion channels and scramblases. TMEM16A and TMEM16F channel activation requires both intracellular Ca^2+^_i_ and membrane depolarization (Pedemonte and Galietta, 2014). We first tested whether pH_i_ has any effect on their voltage-dependent activation of the two channels. We have shown that the *G-pH*_*i*_ plots are parallel over a wide range of activation voltages under given Ca^2+^_i_ (Fig. 1C and 1F), suggesting that pH_i_ is likely to have minimal effect on the voltage-dependent activation of TMEM16A and TMEM16F channels. To further support this observation, we utilized the gain-of-function mutations in the pore-gate domain, namely TMEM16A-Q645A, TMEM16A-L543Q (Fig. 3A), TMEM16F-F518K and TMEM16F-Y563K (Fig. 3D) (Peters *et al*., 2018; T. Le *et al*., 2019). As shown in Fig. 3B and 3E, all these GOF mutations can be activated by membrane depolarization in the absence of Ca^2+^_I_, which allows us to explicitly dissect out the effects of pH_i_ on voltage dependent activation in zero Ca^2+^_i_ (Fig. 3B). When voltage is the only stimulus, pH_i_ effects on the TMEM16A (Fig. 3B-C) and TMEM16F mutant channels (Fig. 3E-F) are almost entirely abolished, as evidenced by the nearly flat *I-pH*_*i*_ curves for all the GOF mutations (Fig.3C and 3F). Based on these results, we conclude that pH_i_ exerts no obvious effect on voltage-dependent activation of TMEM16A and TMEM16F ion channels.

**Figure 3.**
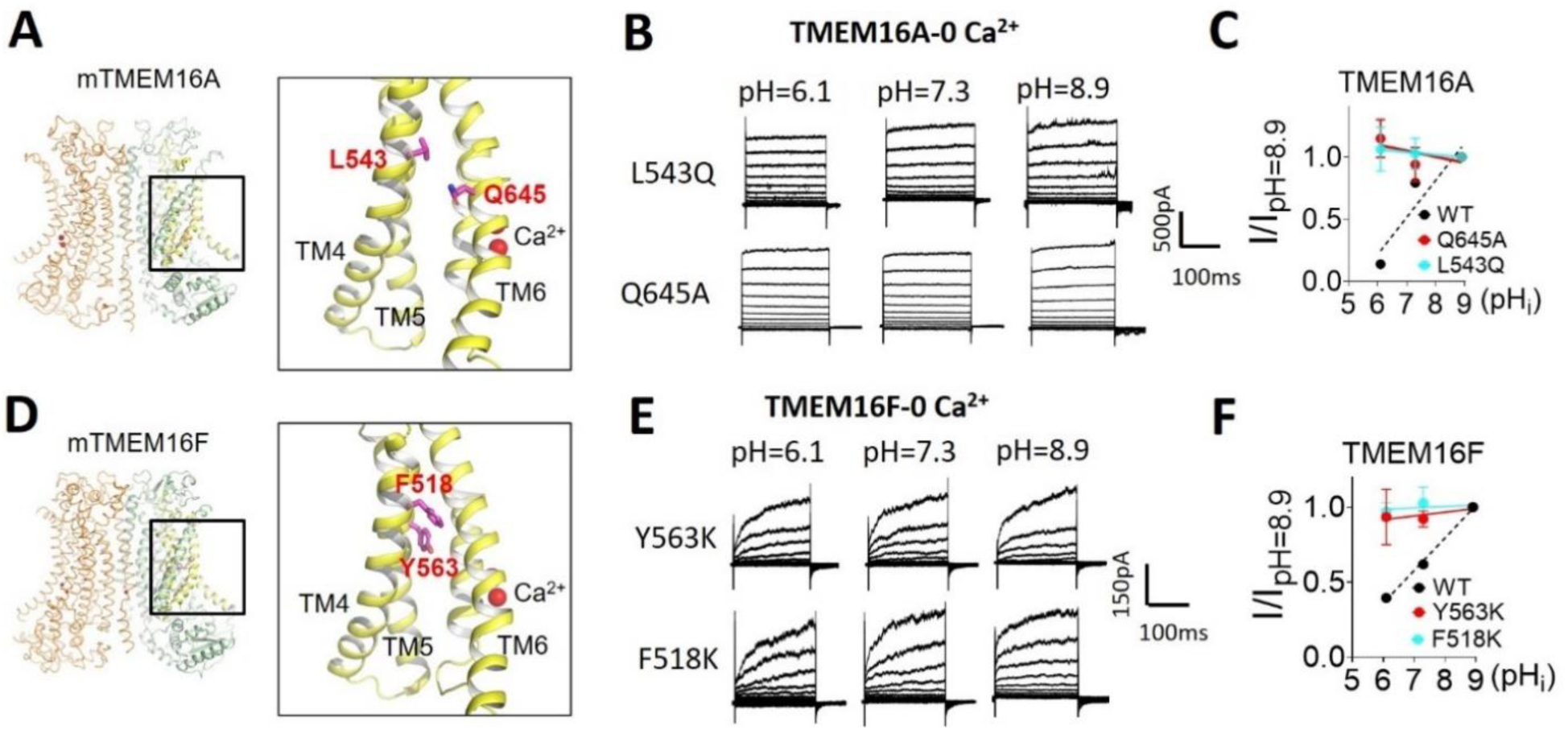
pH_i_ has no effect on TMEM16A or TMEM16F when Ca^2+^ is absent. **(A)** Locations of L543 and Q645 on the TMEM16A structure (PDB 5OYB). **(B)** Representative TMEM16A-L543Q and TMEM16A-Q645A currents recorded from inside-out patches perfused with intracellular solutions containing 0 Ca^2+^ at different pH_i_. **(C)** *I-pH*_*i*_ curve of TMEM16A mutations L543Q and Q645A at 100 mV. Slopes from linear fit for L543Q and Q645A are −0.02 and −0.04, respectively. The *G-pH*_*i*_ curve of WT under 0.5 µM Ca^2^ was replotted as the black dash line. Error bars represent standard deviation (n=7). **(D)** Locations of Y563 and F518 on the TMEM16F structure (PDB 6QP6). **(E)** Representative TMEM16F-Y563K and TMEM16F-F518K currents recorded from inside-out patches perfused with intracellular solutions containing 0 Ca^2+^ at different pH_i_. **(F)** The *I-pH*_*i*_ relationship of TMEM16F mutations Y563K and F518K at 100 mV. Slopes from linear fit for Y563K and F518K are 0.03 and 0.01, respectively. *G-pH*_*i*_ curve of WT under 100 µM Ca^2+^ was replotted as black dash line. Error bars represent standard deviation (n=5). Note that the gain-of-function mutations do not have obvious tail current under 0 Ca^2+^, thereby *I-pHi* relations not *G-pH*_*i*_ were plotted to evaluate their pH_i_ sensitivities.

### pH_i_ regulation of TMEM16F ion channel activity is Ca^2+^ dependent

As lowering pH_i_ has been reported to inhibit Ca^2+^-dependent activation of TMEM16A (Chun *et al*., 2015), we next examined whether pH_i_ effect on TMEM16F channel activation is also through influencing its Ca^2+^ dependence by comparing the pH_i_ effects under 5, 100 and 1000 µM Ca^2+^_i_. Under 5 µM Ca^2+^_i_, TMEM16F channel activation is strongly pH_i_ dependent (Fig. 4A and 4C). However, the pH_i_ dependence under 5 µM Ca^2+^_i_ is apparently different from that under 100 µM Ca^2+^_i_ (Fig. 1D-E): no TMEM16F channel activity can be observed at low pH_i_ (=6.1), and the threshold voltage to activate TMEM16F current is at higher voltage (∼80mV) than the threshold activation voltage under 100 µM Ca^2+^_i_ (Fig.3, A and C). The apparent pH_i_ sensitivity at 100 mV under 5 µM Ca^2+^_i_ increases to 0.32 compared to the pH_i_ sensitivity of 0.20 under 100 µM Ca^2+^_i_, suggesting that pH_i_ sensitivity is enhanced when Ca^2+^_i_ is low (Fig. 4E). On the contrary, the pH_i_ effect is almost completely diminished when the Ca^2+^_i_ is increased to 1000 µM, as seen by the robust and almost identical activation of TMEM16F current from pH_i_ 6.1 to 8.9 and nearly flat *G-V* relationships (Fig. 4B and 4D). Consistently, when saturating Ca^2+^_i_ (100 µM) was applied to TMEM16A-CaCC at different pH_i_, the pH_i_ effect on TMEM16A also diminished (Fig. S2). Our results thus demonstrate that the pH_i_ regulation of TMEM16F and TMEM16A channel is highly Ca^2+^ dependent in which saturating Ca^2+^_i_ (1000 µM for TMEM16F and 100 µM for TMEM16A) eliminates their pH_i_ sensitivity, whereas lower Ca^2+^_i_ boosts pH_i_ sensitivity.

**Figure 4.**
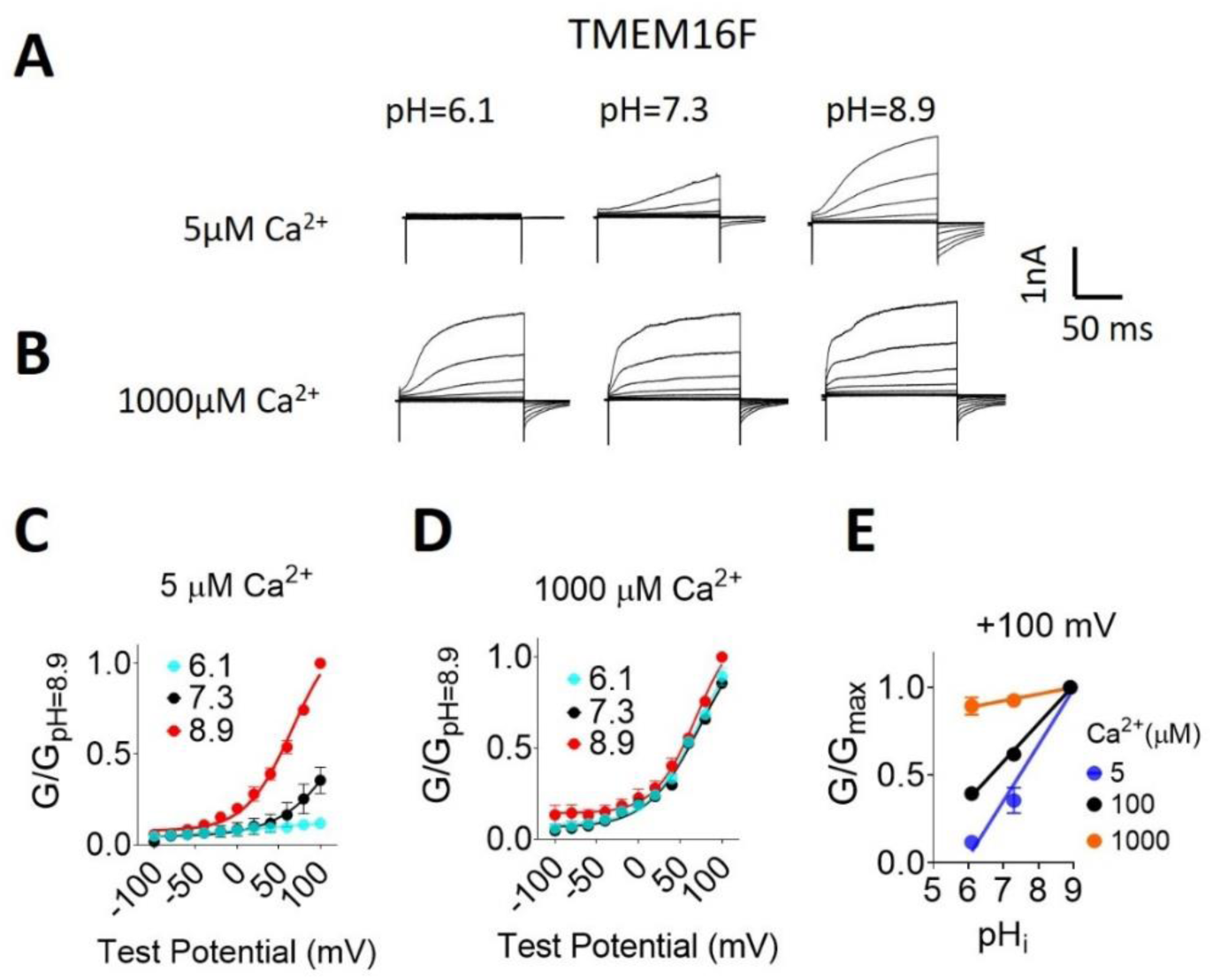
pH_i_ regulation of TMEM16F channel activity is Ca^2+^ dependent. **(A), (B)** Representative TMEM16F currents recorded from inside-out patches perfused with intracellular solutions containing 5 µM and 1000 µM Ca^2+^, respectively. All traces shown in each panel were from the same patch. Currents were elicited by voltage steps from −100 to +100 mV with 20 mV increments. The holding potential was −60 mV. **(C), (D)** Mean *G-V* relations of the TMEM16F currents from (A) and (B), respectively. Relative conductance was determined by measuring the amplitude of tail currents 400 μs after repolarization to a fixed membrane potential (−60 mV). The smooth curves represent Boltzmann fits. Error bars represent SEM (n = 5). **(E)** pH_i_ sensitivity of TMEM16F current at +100 mV was measured by the slope of the the *G-pH*_*i*_ relationship. Mean conductance at different Ca^2+^ concentration was normalized to the maximum conductance at pH_i_=8.9 and +100 mV. Averaged slopes from linear fit for 5 and 1000 µM Ca^2+^ are 0.32 and 0.04, respectively. The *G-pH*_*i*_ curve at 100 µM Ca^2+^ was replotted as black line for reference. Error bars represent standard deviation (n=5).

### pH_i_ regulation of TMEM16F lipid scrambling activity is Ca^2+^ dependent

We next addressed whether the pH_i_ regulation on TMEM16F lipid scrambling is also Ca^2+^_i_ dependent using our lipid scrambling fluorometry assay (Fig. 2A). Under 5 µM Ca^2+^_i_, TMEM16F lipid scrambling is strongly pH_i_ dependent (Fig. 5A-B and Movie S3). However, comparing with 100 µM Ca^2+^_i_, t_1/2_ under 5 µM Ca^2+^_i_ is significantly delayed at pH_i_ of 7.3 and 8.9 (Fig. 5C and Fig. S3A). In addition, the maximum lipid scrambling rates as quantified by slope k are significantly reduced under 5 µM Ca^2+^_i_ comparted to 100 µM Ca^2+^_i_ (Fig. 5D and Fig. S3B). TMEM16F scrambling activity under 5 µM Ca^2+^_i_ is completely abolished when pH_i_ was 6.1. This is distinct from TMEM16F scrambling in 100 µM Ca^2+^_i_ and pH_i_ 6.1, under which condition TMEM16F-mediated lipid scrambling can be clearly observed (Fig. 2B-D). On the contrary, when saturated intracellular Ca^2+^ (1000 µM) was applied, fast and robust PS exposure were detected at all intracellular pH_i_ (Fig. 5E and Movie S4). In addition, the t_1/2_ and the maximum lipid scrambling rates (slope k) of TMEM16F lipid scrambling are comparable regardless of pH_i_ (Fig. 5G-H and Fig. S3). Our imaging experiments thus demonstrate that similar to TMEM16F channel activity (Fig. 4), pH_i_ regulation of TMEM16 CaPLSase activity is also highly Ca^2+^_i_ dependent (Fig. S3).

**Figure 5.**
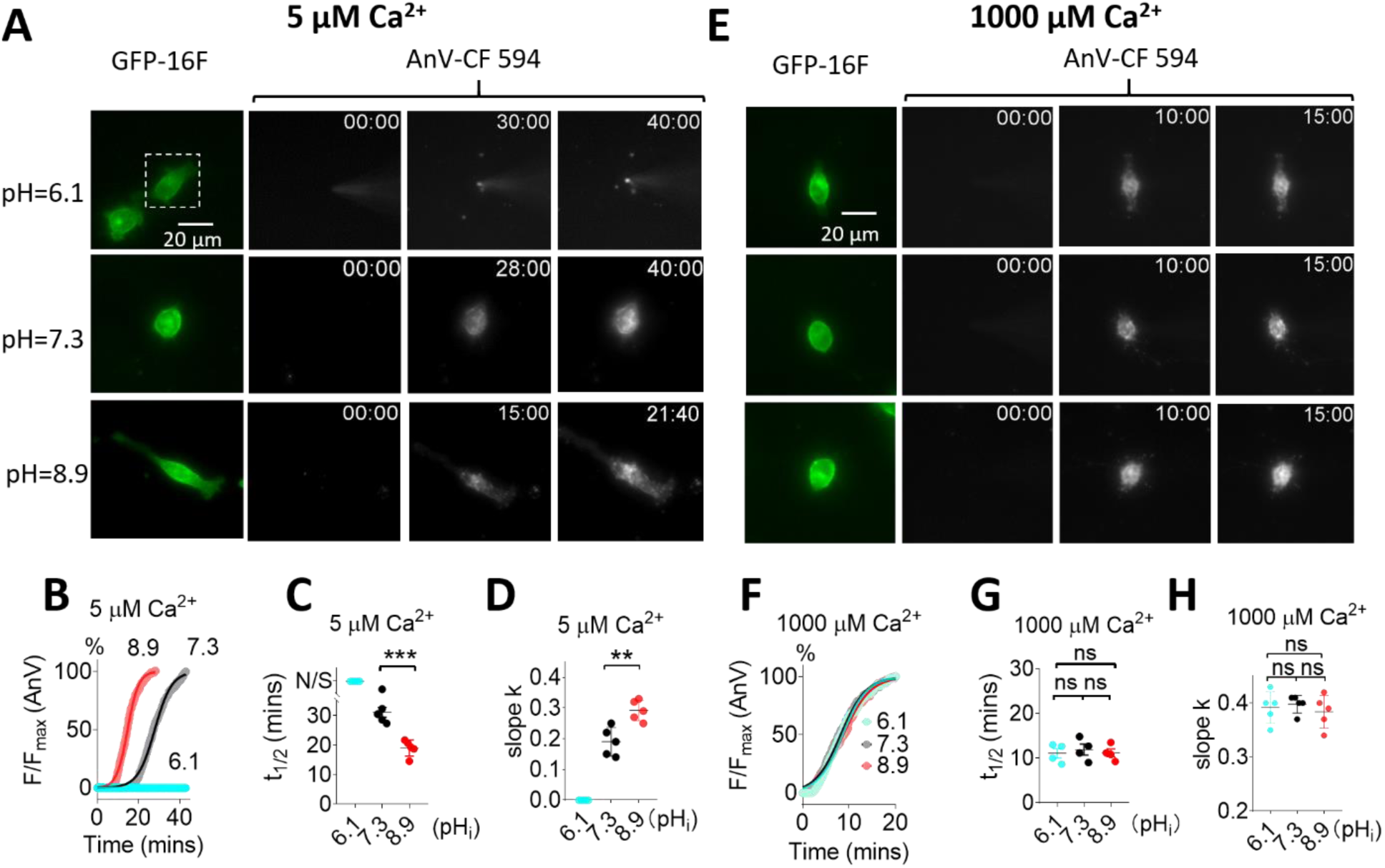
pH_i_ regulation of TMEM16F scrambling activity is Ca^2+^ dependent. **(A)** Representative images of TMEM16F-eGFP scrambling activity under 5 μM intracellular Ca^2+^ with different pH_i_. The white dotted rectangles in the top row demarcate the patch clamped TMEM16F-eGFP expressing cells. For the AnV-CF 594 signals on the right, the first column is fluorescence signal immediately after forming whole-cell configuration; the second column is the time point (minutes followed by seconds, top right corner) when fluorescence intensity reaches half maximum (t_1/2_) and the last column is the time point when fluorescence reaches plateau. No obvious AnV-CF 594 signal can be detected in pH_i_ 6.1 over 40 minutes. **(B)** The time courses of the AnV fluorescent intensity for TMEM16F activated by 5 μM Ca^2+^ under different pH_i_ in (A). The smooth curves represent fits to the logistic equation similar to Fig.2B. **(C)** The t_1/2_ at different pH_i_ under 5 μM Ca^2+^. N/S at pH_i_=6.1 represents no scrambling. Error bars represent SEM (n = 5). **(D)** The slopes at different pH_i_ under 5 μM Ca^2+^. **(E)** Representative images of TMEM16F-eGFP scrambling activity under 1000 μM intracellular Ca^2+^ with different pH_i_. For the AnV-CF 594 signal on the right, the first column is fluorescence signal immediately after forming whole-cell configuration; the second column is the time point when fluorescence intensity reaches half maximum (t_1/2_) and the last column is the time point when fluorescence reaches plateau. **(F)** The time courses of the AnV fluorescent intensity for TMEM16F activated by 1000 μM Ca^2+^ under different pH_i_ in (E). The smooth curves represent fits to the logistic equation. **(G)** The t_1/2_ of lipid scrambling at different pH_i_ under 1000 μM Ca^2+^. **(H)** The slopes k at different pH_i_ under 1000 μM Ca^2+^. Error bars represent SEM (n = 5). Statistics were done using one-way ANOVA followed by Tukey’s test. **=p<0.01, ***=p < 0.001 and ns (not significant): p>0.05.

### pH_i_ alters Ca^2+^ sensitivity of TMEM16F ion channel

Having shown that pH_i_ regulation of TMEM16F is Ca^2+^_i_ dependent but not voltage dependent, we next tested the hypothesis that pH_i_ may directly influence the Ca^2+^ binding sites of TMEM16F following the same pH_i_ regulation mechanism of TMEM16A. If this is the case, protonation and deprotonation of the Ca^2+^ binding residues would reduce or increase the Ca^2+^ sensitivity of TMEM16F, respectively. In order to test this hypothesis, we measured the apparent Ca^2+^ sensitivity of TMEM16F under different pH_i_ using inside-out patches (Fig. 6A). Our results showed that the EC_50_ of TMEM16F Ca^2+^-dose response under physiological pH_i_ is 6.2±2.2 µM (Fig. 6B-C), comparable with previous studies (Yang *et al*., 2012). When pH_i_ drops to 6.1, TMEM16F Ca^2+^ sensitivity decreases more than 20 folds (EC_50_ = 144.1±6.8 µM). In stark contrast, when pH_i_ is switched to 8.9, TMEM16F Ca^2+^ sensitivity is dramatically enhanced (EC_50_ = 1.2±1.2 µM). These results thus further support that pH_i_ works on the Ca^2+^ binding sites to exert its regulatory effects.

**Figure 6.**
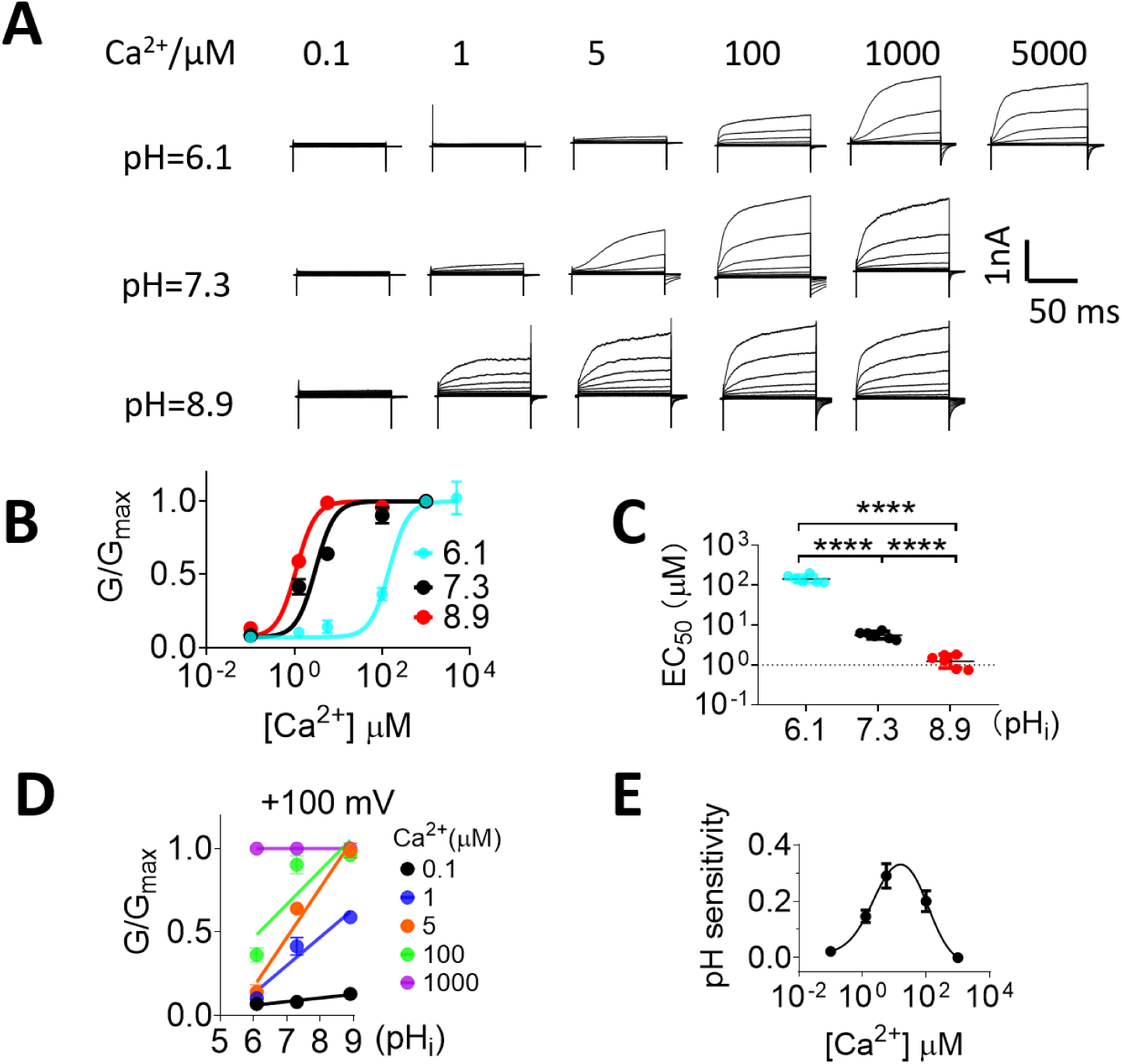
pH_i_ alters Ca^2+^ binding affinity of TMEM16F. **(A)** Representative TMEM16F currents recorded from inside-out patches perfused with intracellular solutions containing 0.1,1,5,100,1000 and 5000 (pH=6.1 only) µM Ca^2+^ at different pH_i_. Currents were elicited by voltage steps from −100 to +100 mV with 20 mV increments. The holding potential was −60 mV. **(B)** Ca^2+^ dose-response of mTMEM16F channel at +100 mV with different pH_i_. The smooth curves represent the fits to the Hill equation: *G/G*_*max*_*=G*_*1000*_*/(1+(K*_*D*_*/[Ca*^*2+*^*])H)*, wherein *K*_*D*_ is the apparent dissociation constant, *H* is the Hill coefficient, and *G*_*1000*_ is the conductance with 1000 µM Ca^2+^ at given pH_i_. The error bars represent SEM (n=5). **(C)** Half maximal effective concentrations of Ca^2+^ (EC_50_) at pH_i_ 6.1, 7.3 and 8.9 were 144.1±6.8, 6.2±2.2 and 1.2±1.2 µM, respectively. p-values were determined with Tukey test comparisons following one-way ANOVA. **** p < 0.0001. **(D)** The *G-pH*_*i*_ relationship of TMEM16F current at +100 mV under different Ca^2+^ concentrations. Solid lines represent linear fits. **(E)** The relationship of pH_i_ sensitivity and intracellular Ca^2+^ concentration. The pH_i_ sensitivity values were slopes from linear fit shown in (D) under different Ca^2+^ concentrations. The smooth line was fitted with bell-shaped dose response curve using Graphpad Prism with peak pH sensitivity of 0.33 at ∼15 µM Ca^2+^.

Next, we extracted the relative conductance (*G/G*_*Max*_) from Fig. 6B and plotted the *G*-pH_i_ relationship for each Ca^2+^_i_ (Fig. 6D). The *G*-pH_i_ relationships under different Ca^2+^_i_ are not parallel, which is in stark contrast to the parallel *G*-pH_i_ relationships under different membrane voltages (Fig. 1F). As the slope of the *G*-pH_i_ relationship represents the apparent pH_i_ sensitivity, we constructed the pH_i_ sensitivity-Ca^2+^_i_ relationship curves in Fig. 6E. Interestingly, the pH_i_ sensitivity-Ca^2+^_i_ curve displays a bell-shaped distribution, which peaks around 15 µM Ca^2+^_i_. Recapitulating the bell-shaped Ca^2+^response curves of lns(1,4,5)P_3_ (IP_3_)-and Ca^2+^-gated IP_3_R and ryanodine receptor (RyR) channels (Bezprozvanny, Watras and Ehrlich, 1991), the bell-shaped pH_i_ sensitivity-Ca^2+^_i_ curve of TMEM16F demonstrates that TMEM16F is strictly pH_i_ sensitive under physiological range of Ca^2+^_i_. TMEM16F channel activation becomes more and more sensitive to pH_i_ when Ca^2+^_i_ elevates from resting level of 0.1 µM until its pH_i_ sensitivity peaks around 15 µM Ca^2+^_i_. When Ca^2+^_i_ increases beyond 15 µM Ca^2+^_i_, TMEM16F pH_i_ sensitivity sharply decreases. Our analysis thus defines the physiological range of TMEM16F pH_i_ regulation.

### Partially disrupting the Ca^2+^ binding sites enhances pH_i_ sensitivity

To further prove that pH_i_ directly affects Ca^2+^ binding residues, we examine the pH_i_ sensitivity of E667Q, the Ca^2+^ binding site mutation that markedly reduces TMEM16F Ca^2+^ sensitivity with EC_50_ of 2.8 mM (Yang *et al*., 2012) (Fig. 7 A). In stark contrast to the lack of pH_i_ sensitivity of wildtype (WT) TMEM16F under 1000 µM Ca^2+^_i_ (Fig. 4B, 4D and 4E), the activation of the loss-of-function (LOF) E667Q becomes strongly pH_i_ dependent in 1000 µM Ca^2+^_i_ (Fig. 7B-D). Similarly, another Ca^2+^ binding site LOF mutation E670Q also exhibits markedly enhanced pH_i_ sensitivity under 1000 µM Ca^2+^_i_ (Fig. S4). As a control, we examined the pH_i_ sensitivity of Q559K, a mutation of the pore lining residue Q559 that affects TMEM16F ion selectivity without obvious effect on Ca^2+^ binding (Fig. 7A) (Yang *et al*., 2012; Ye *et al*., 2019). As shown in Fig. 7E-G, the pH_i_ sensitivity of Q559K under 100 µM Ca^2+^_i_ is almost identical with the pH_i_ sensitivity of WT TMEM16F channel. Consistent with what we observed in TMEM16F, the LOF mutation of a TMEM16A Ca^2+^ binding residue E730Q also renders the CaCC with strong pH_i_ sensitivity under 100 µM Ca^2+^_i_ compared with the lack of pH_i_ sensitivity of WT TMEM16A under this saturating Ca^2+^_i_ (Fig. S2 and Fig. S5). Taken together, our systematic biophysical characterizations and mutagenesis experiments explicitly illustrate a pH_i_ regulatory mechanism for both TMEM16F and TMEM16A. According to this mechanism, pH_i_ regulates the activation of these TMEM16 proteins through protonation and deprotonation of their Ca^2+^ binding sites, which in turn reduces and enhances their Ca^2+^ binding affinity, respectively (Fig. 7G). As the Ca^2+^ binding residues are highly conserved, this pH_i_ regulatory mechanism may also apply to other TMEM16 family members.

**Figure 7.**
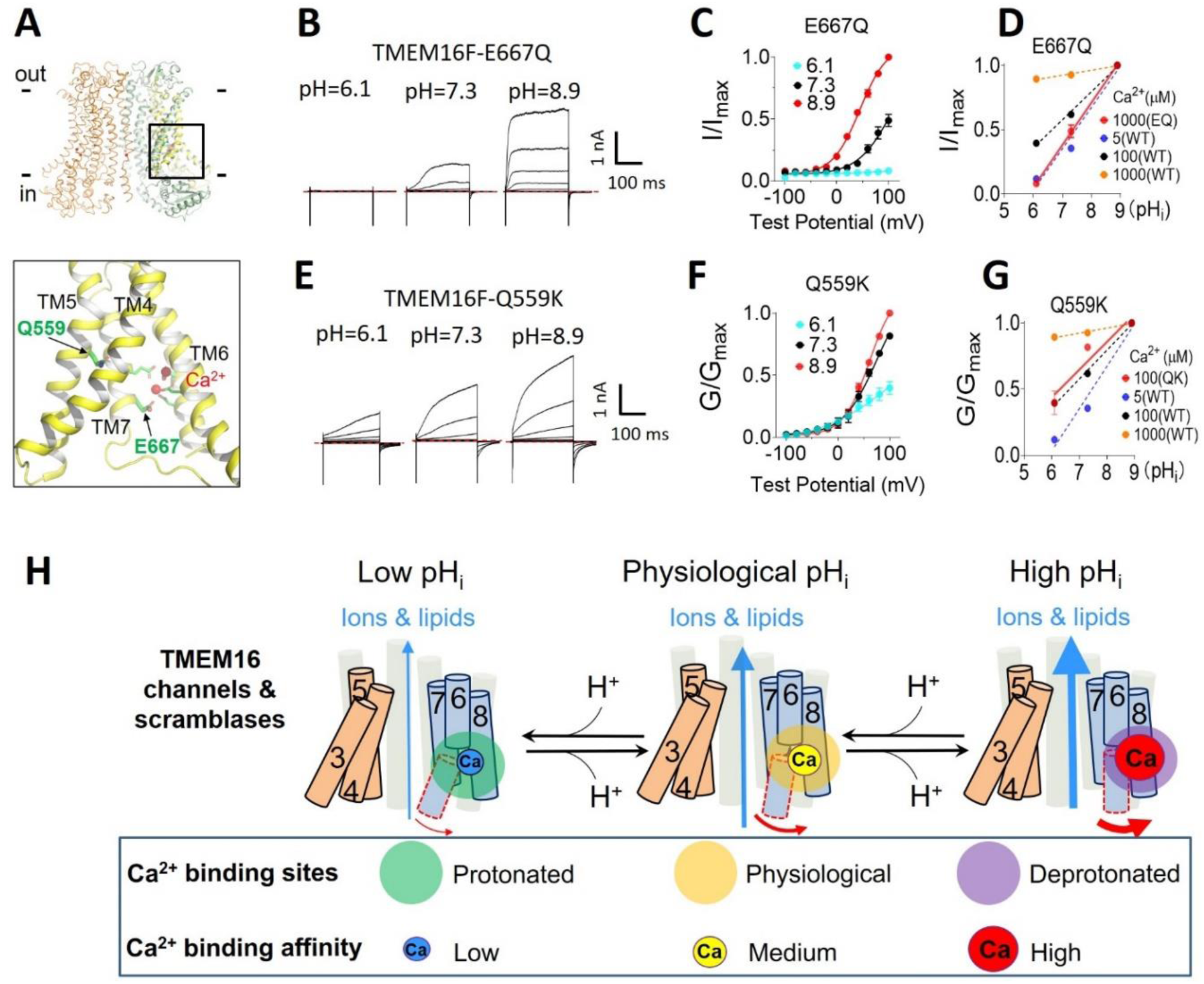
pHi specifically works on the Ca^2+^ binding sites of TMEM16F to regulate its activities. **(A)** Locations of the pore-lining residue Q559 and the Ca^2+^ binding residue E667 on the TMEM16F structure (PDB ID: 6QP6). **(B)** Representative TMEM16F-E667Q currents recorded from inside-out patches perfused with intracellular solutions containing 1000 µM Ca^2+^ at different pH_i_. **(C)** The *I-V* curve of E667Q currents at different pH_i_. **(D)** The pH_i_ sensitivity of E667Q (EQ) evaluated by the *I-pH*_*i*_ relationship. The slope of the *I-pH*_*i*_ relationship for E667Q at 1000 µM Ca^2+^ is 0.30±0.02 shown in red solid line. The *I-pH*_*i*_ curves of WT at different Ca^2+^ concentrations were plotted as dashed lines as references. **(E)** Representative TMEM16F-Q559K currents recorded from inside-out patches perfused with intracellular solutions containing 100 uM Ca^2+^ at different pH_i_. **(F)** The *G-V* curves of Q559K currents at different pH_i_. **(G)** The pH_i_ sensitivity of Q559K (QK) evaluated by the *G-pH*_*i*_ relationship. The slope of the *G-pH*_*i*_ relationship for Q559K at 100 µM Ca^2+^ is 0.20±0.02 shown in red solid line. The *G-pH*_*i*_ curves of WT at different Ca^2+^ concentrations were also plotted as dashed lines as references. (**H)** A cartoon illustration of pH_i_ regulation of TMEM16 channels and scramblases. Under low pH_i_ condition, protonation of Ca^2+^ binding residues can significantly reduce Ca^2+^ binding affinity of TMEM16 proteins and suppress their activation. In contrast, high pH_i_ deprotonates the Ca^2+^ binding residues, enhances Ca^2+^ binding and TMEM16 activation. The size of arrows represents the open probability of TMEM16 proteins. The size of Ca^2+^ ions represents the strength of apparent Ca^2+^ binding affinity.

### Intracellular alkalization promotes TMEM16F activities in a cancer cell line

Given that intracellular alkalization is one of the hallmarks of cancer cells (Cardone, Casavola and Reshkin, 2005; White, Grillo-Hill and Barber, 2017) and TMEM16F is highly expressed in a wide spectrum of tumors (Jacobsen *et al*., 2013; Wang *et al*., 2018; Xuan, Wang and Xie, 2019), we next sought to examine whether endogenous TMEM16F in tumor cells can also be strongly promoted by intracellular alkalization as we observed in the HEK293 cells exogenously expressed with TMEM16F. To address this, we utilized a human choriocarcinoma cell line BeWo that highly expresses TMEM16F CaPLSases to enable cell-cell fusion of the placental trophoblast tumor cells (Zhang *et al*., 2020). As shown in Fig. 8, mild intracellular alkalization from pH_i_ 7.2 to 7.7, the pH_i_ range observed in many cancer cells (Webb *et al*., 2011) can significantly enhance both TMEM16F ion channel (Fig. 8A-C) and CaPLSase activities (Fig. 8D-G and Movie S5). In contrast, intracellular alkalization to pH_i_ 7.7 is incapable of inducing any TMEM16F-like current or CaPLSase activity in our TMEM16F deficient (KO) BeWo cells (Zhang *et al*., 2020) (Fig. S6 and Movie S6). Our characterizations of the endogenous TMEM16F in BeWo cancer cell line thus demonstrate that intracellular alkalization indeed enhances endogenous TMEM16F activities in tumor cells.

**Figure 8.**
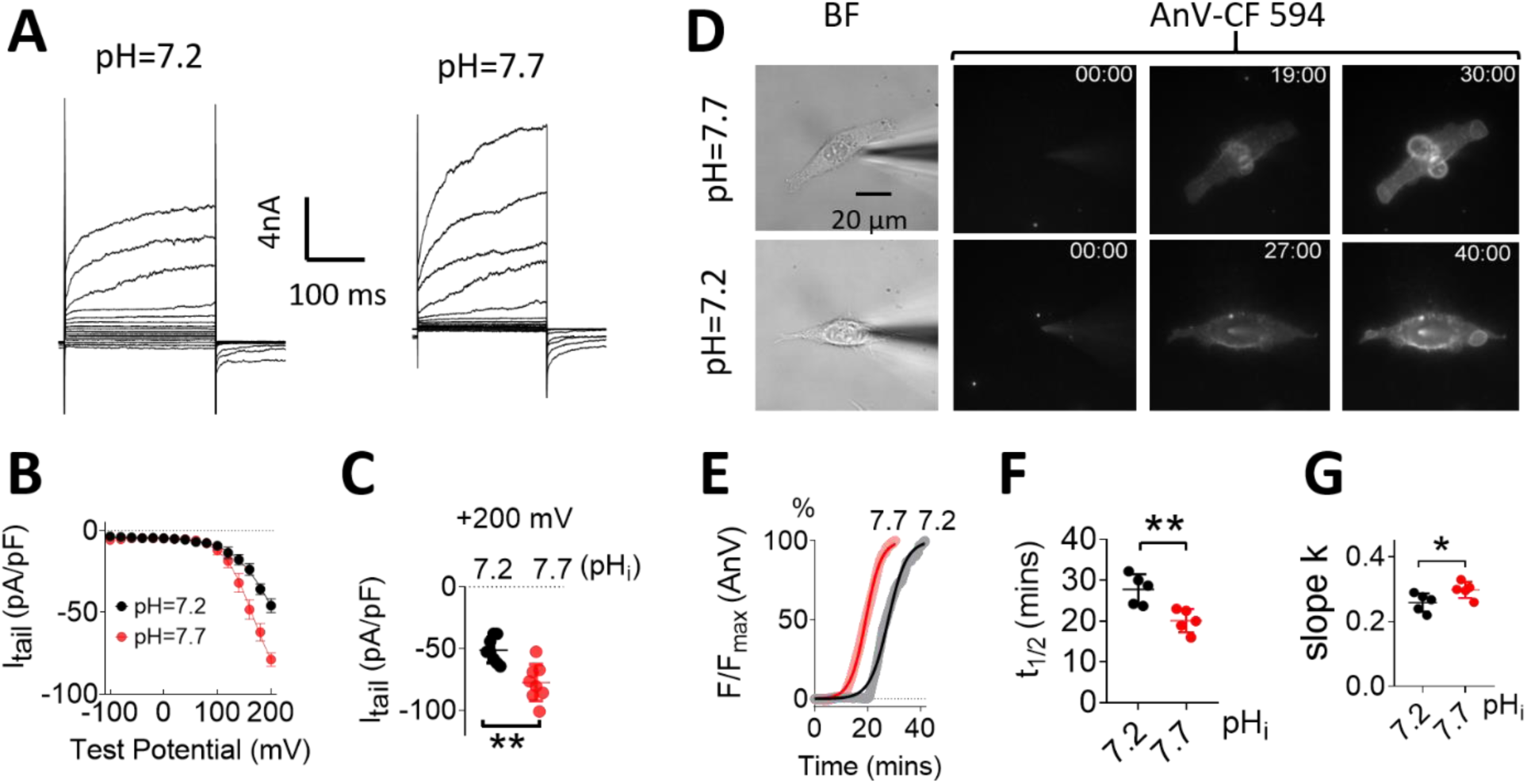
Intracellular alkalization potentiates the activation of endogenous TMEM16F CaPLSase in choriocarcinoma BeWo cells. **(A)** Representative whole-cell currents recorded from wildtype (WT) BeWo cells perfused with intracellular solutions containing 5 µM Ca^2+^ at different pH_i_ (7.2 and 7.7). **(B)** Mean *I-V* relation using tail currents at different pH_i_. Error bars represent SEM (n=5). **(C)** Comparison of the tail currents at +200 mV under different pH_i_. Error bar represents SEM (n=5). **(D)** Representative CaPLSase activity of WT BeWo cells at different pH_i_. BF, bright field. For AnV-CF 594 signals, the first column is recorded immediately after forming whole-cell patch configuration; the second column is the time point when fluorescence intensity reaches half maximum (t_1/2_) and the last column is the time point when fluorescence reaches plateau. The time on the top right corner shows minutes followed by seconds. **(E)** The time course of endogenous CaPLSases under pH_i_ 7.2 and 7.7. The fluorescence signal was normalized to its maximum intensity for each pH_i_. The smooth curves represent fits to the logistic equation. **(F)** t_1/2_ at different pH_i_. Averaged t_1/2_ for pH_i_=7.2 and 7.7 are 27.7±3.3 minutes (n=5) and 20.1±2.5 minutes (n=5), respectively. (G) The slopes at different pH_i_. Averaged slopes for pH_i_=7.2 and 7.7 are 0.26±0.03 (n=5) and 0.30±0.02 (n=5), respectively. Values represent mean ± SEM and statistics were done using one-way ANOVA followed by Tukey’s test. * =p<0.05 and ** =p<0.01.

## Discussion

In this study, we report that pH_i_ can effectively regulate TMEM16F CaPLSase and ion channel activities. Through systematic investigation of TMEM16F and comparison with TMEM16A-CaCC that is inhibited by intracellular protonation, we found that pH_i_ regulates TMEM16F and TMEM16A activation through the same molecular mechanism. Our biophysical characterizations and mutagenesis studies explicitly show that the Ca^2+^ binding residues within the pore-gate domains of the TMEM16 proteins serve as the pHi sensors (Fig. 7H). Protonation of the carboxylate groups of the Ca^2+^ binding residues prevents Ca^2+^ binding, thereby hindering TMEM16 activation (Fig. 6C). On the contrary, deprotonation of the carboxylate groups of the Ca^2+^ binding residues facilitates Ca^2+^ binding, thereby promoting TMEM16 activation.

Histidine, the most titratable amino acid under physiological pH_i_ range, is unlikely to be the pH_i_ sensor for TMEM16F activation. The previous mutagenesis study of TMEM16A demonstrates that all the intracellular histidine to alanine mutations do not alter the inhibitory effect of proton on TMEM16A activation (Chun *et al*., 2015). As some of these histidine residues are conserved between TMEM16F and TMEM16A, it is thus plausible to assert that the equivalent histidine residues also do not contribute to TMEM16F pH_i_ sensing. In addition, our characterizations of the GOF mutations and LOF mutations in this study further support that the intracellular histidine residues are unlikely to involve in pH_i_ regulation. For the GOF mutations shown in Fig. 3, pH_i_ losses its effects on TMEM16F and TMEM16A activation in the absence of Ca^2+^_i_. If some of the histidine residues contribute to pH_i_ sensing, we would still be able to see some residual pH_i_ sensitivity. In addition, the dramatically enhanced pH_i_ sensitivity for the LOF mutations of the Ca^2+^-binding sites in TMEM16F (Figs. 7B-C and S4) and TMEM16A (Fig. S5) further support that the Ca^2+^ binding resides but not the intracellular histidine residues are responsible for pH_i_ sensing.

Interestingly, pH_i_ sensitivity of TMEM16F is highly Ca^2+^ dependent and exhibits a bell-shaped relationship with Ca^2+^_i_ (Fig. 6E). According to this relationship, saturating Ca^2+^_i_can override the pH_i_ effects on the protonation states of the Ca^2+^ binding residues, thereby eliminates pH_i_ sensitivity for WT TMEM16F and TMEM16A (Figs. 4E, 5F and Fig. S2). Mutating the Ca^2+^ binding residues dramatically reduces TMEM16F and TMEM16A apparent Ca^2+^ sensitivities (Yang *et al*., 2012; Yu *et al*., 2012; Tien *et al*., 2014; Alvadia *et al*., 2019). This disrupts the balance between pH_i_ and Ca^2+^_i_ so that the protonation states of Ca^2+^ binding site in these LOF mutant proteins gain more weight on influencing Ca^2+^ binding affinity. Therefore, the Ca^2+^ binding site mutations become pH_i_ sensitivity under high Ca^2+^_i_, which are saturating for WT TMEM16 proteins (Fig. 7D, Figs. S4D and S5D). On the other hand, when Ca^2+^_i_ is very low and voltage plays a more prominent role in activating the TMEM16 proteins, the pH_i_ sensitivities of TMEM16F and TMEM16A reduces. This is likely because pH_i_ has negligible impact on voltage-dependent activation of these proteins as evidenced by the near parallel *G-pH*_*i*_ relationships across wide range of activation voltages (Fig. 1C and 1F), as well as the lack of pH_i_ effects on the voltage dependent activation of the GOF mutations of TMEM16F and TMEM16F in the absence of Ca^2+^_i_ (Fig. 3). For TMEM16F, the pH_i_ sensitivity gradually increases from resting 0.1 µM Ca^2+^ to its maximum sensitivity at ∼15 µM Ca^2+^_i_. It is worth noting that all our experiments were conducted at room temperature. According to a recent publication (Lin *et al*., 2019), TMEM16F is more sensitive to Ca^2+^_i_ at 37°Ctemperature.. It is therefore plausible that pH_i_ may be more sensitive to Ca^2+^_i_ under physiological

Intracellular alkalization is one of the hallmarks of cancer cells (Webb *et al*., 2011). A number of distinct ion transporters and pumps, including the Na^+^-H^+^ exchangers (Lauritzen *et al*., 2012), the H^+^/K^+^-ATPase proton pump (Goh, Sleptsova-Freidrich and Petrovic, 2014) and the Na^+^-driven bicarbonate transporters (McIntyre *et al*., 2016), have been known to contribute to intracellular alkalization. Different from the extensive understanding of pH_i_ dysregulation in cancer cells, how intracellular alkalization affects cancer cell function is still elusive. Here we show that a mere 0.5 pH_i_ unit increase from the physiological pH_i_ of 7.2 can significantly enhance both TMEM16F CaPLSase and ion channel activation at room temperature. Thus, the dual functional TMEM16F CaPLSase and ion channel can be a new pH_i_ sensing protein that responds to intracellular alkalization.

TMEM16F has been identified in a wide variety of cancer cells; and according to the Human Protein Atlas (http://www.proteinatlas.org), the high expression level of TMEM16F is associated with the overall prognosis of a number cancers including breast and cervical cancers. Although it is unclear how TMEM16F contributes to tumor growth and cancer progression, it has been reported that genetic manipulations of TMEM16F can change cancer cell proliferation and migration (Jacobsen *et al*., 2013; Wang *et al*., 2018; Xuan, Wang and Xie, 2019). Interestingly, loss of membrane phospholipid asymmetry is a salient feature of many cancerous cells, whose cell surfaces display an elevated amount of PS (Riedl *et al*., 2011; Zhao *et al*., 2011). It is still unclear which phospholipid transporters are responsible for the enhanced PS exposure on the surfaces of the cancer cells. Nevertheless, no sign of apoptosis (Fidler *et al*., 1991) and the beneficial effects of PS exposure to cancer cell survival (Schröder-Borm, Bakalova and Andrä, 2005; He *et al*., 2009; Kenis and Reutelingsperger, 2009; Gerber *et al*., 2011; Blanco *et al*., 2014; Zhang *et al*., 2014) suggest that CaPLSases instead of caspase-dependent lipid scramblases may play important roles in facilitating PS exposure in cancer cells. We show in this study that the CaPLSase and ion channel activities of the endogenous TMEM16F in a human choriocarcinoma cell line indeed can be pronouncedly promoted by intracellular alkalization (Fig. 8). Our current investigation of the mechanism of TMEM16F pH_i_ regulation thus lays a foundation to further understand the role of TMEM16F in cancer and other physiological or pathological conditions, in which pH_i_ fluctuates or is dysregulated. The shared molecular mechanism of pH_i_ regulation between TMEM16F and TMEM16A identified in this study will also facilitate our understanding the regulatory mechanism and physiological functions of other TMEM16 family members.

## Methods and Materials

### Cell lines and culture

HEK293T and BeWo cells are authenticated by Duke Cell Culture Facility. HEK293T cell line with stable expression of C-terminally eGFP-tagged mTMEM16F was a gift from Dr. Min Li. The TMEM16F and BeWo deficient (KO) HEK293T cell line were generated using CRISPR-Cas9 and have been validated in our recent studies (Le, Le and Yang, 2019; T. Le *et al*., 2019; Zhang *et al*., 2020). All our mutagenesis studies were conducted in the TMEM16F-KO HEK293T cells. HEK293T cells were cultured with Dulbecco’s modified Eagle’s medium (DMEM; Gibco, #11995-065) supplemented with 10% fetal bovine serum (Sigma, cat. F2442) and 1% penicillin– streptomycin (Gibco, #15-140-122). BeWo cells were cultured in Dulbecco’s Modified Eagle Medium-Hams F12 (DMEM/F12) medium (Gibco, #11320-033), supplemented with 10% FBS and 1% penicillin/streptomycin. All cells were cultured in a humidified atmosphere with 5% CO2 at 37°C.

### Mutagenesis and transfection

The murine TMEM16F (Open Biosystems cDNA # 6409332) and murine TMEM16A (Open Biosystems cDNA # 30547439) coding sequences are in pEGFP-N1 vector, resulting in eGFP tags on their C-termini. Single point mutations were generated using QuikChange site-directed mutagenesis kit (Agilent) and majorities of them have been reported in previous publications. The plasmids were transiently transfected to the TMEM16F-KO HEK 293T cells by using X-tremeGENE9 transfection reagent (Millipore-Sigma). Cells grown on poly-L-lysine (PLL, Sigma) coated coverslips placed in a 24-well plate. Medium was changed 6 hours post transfection with either regular (Gibco, #11995-065) or Ca^2+^-free DMEM (Gibco, #21068-028). Experiments were proceeded 24–48 hours after transfection.

### Electrophysiology

All currents were recorded in either inside-out or whole-cell configurations 24-48 hours after transfection using an Axopatch 200B amplifier (Molecular Devices) and the pClamp software package (Molecular Devices). The glass pipettes were pulled from borosilicate capillaries (Sutter Instruments) and fire-polished using a microforge (Narishge) to reach resistance of 2–3 MΩ.

For inside-out patch, the pipette solution (external) contained (in mM): 140 NaCl, 10 HEPES, 1 MgCl_2_, adjusted to pH 7.3 (NaOH), and the bath solution contained 140 mM NaCl, 10 mM HEPES, 5 EGTA, adjusted to pH 7.3 (NaOH). Zero Ca^2+^ internal solution contains (in mM): 140 NaCl and 10 HEPES, 5 EGTA. Due to the difficulties to accurately measure and control free Ca^2+^ under different pH_i_, we directly added CaCl_2_ into a solution containing (in mM): 140 NaCl and 10 HEPES in the absence of Ca^2+^ chelator. We may have underestimated the free Ca^2+^ concentrations. But this seems to be the only way to make sure the free Ca^2+^ concentrations keep the same under different pHi. Different pH levels were adjusted either by NaOH or HCl as required.

For whole cell recordings, the pipette solution (internal) contained (in mM): 140 CsCl, 1 MgCl_2_, 10 HEPES, plus CaCl_2_ to obtain the desired free Ca^2+^ concentration. pH was adjusted either by CsOH or HCl as desired. The bath solution contained 140 mM NaCl, 10 mM HEPES, 1 MgCl_2_ and adjusted to pH 7.3 (NaOH). Procedures for solution application were as employed previously (S. C. Le *et al*., 2019). Briefly, a perfusion manifold with 100–200 μm tip was packed with eight PE10 tubes. Each tube was under separate valve control (ALA-VM8, ALA Scientific Instruments), and solution was applied from only one PE10 tube at a time onto the excised patches or whole-cell clamped cells. All experiments were at room temperature (22–25°C). All the chemicals for solution preparation were obtained from Sigma-Aldrich.

### Phospholipid scrambling fluorometry

The phospholipid scrambling fluorometry was adapted from the method developed by the Hartzell laboratory (Yu *et al*., 2015). Instead of delivering and detecting current, the glass pipettes under whole-cell configuration were used to achieve precise control of intracellular pH and Ca^2+^. Briefly, the cells were seeded and transfected on poly-L-lysine (Sigma)-coated coverslips prior to the experiments. Glass pipettes were prepared and filled with internal solution as mentioned previously in the electrophysiology section. After focusing on the cell surface eGFP signal from the expressed TMEM16F-eGFP, the light filter set was switched to detect Annexin V-CF594 signal (594 nm). Annexin V-CF 594 (Biotium # 29011) was diluted at 0.5 µg ml^−1^ with Annexin V-binding solution (10 mM HEPES, 140 mM NaCl, 2.5 mM CaCl_2_, pH 7.4) and then added into the ALA perfusion system as mentioned above. The perfusion manifold was lowered next to the cell to be patched. Once the whole-cell configuration was established, the perfusion valve was simultaneously opened and Anv solution was flushed to the patch-clamped cell. At the same time, image acquisition started with intervals of 5-10 seconds.

### Data Analysis

*G-V* curves were constructed from tail currents measured 200-400 μs after repolarization. In the cases when the tail currents were tiny, the steady-state peak currents were used to build the *I-V* relation. For the *G-V* curves obtained from the same patch, the conductance was normalized to the maximal conductance obtained at pH 8.9 and the highly depolarization voltage. Individual *G-V* curves were fitted with a Boltzmann function,

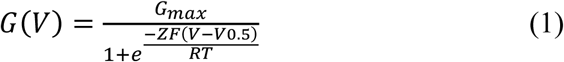

where *G*_*max*_ denotes the fitted value for maximal conductance at a given pH, *V*_*0*.*5*_ denotes the voltage of half maximal activation of conductance, *z* denotes the net charge moved across the membrane during the transition from the closed to the open state and *F* denotes the faraday constant.

Dose response curves were fitted with Hill equation,

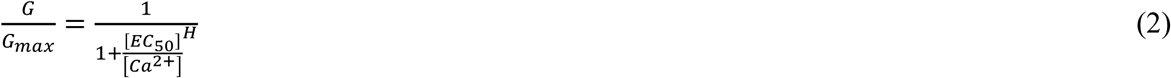

where *G/G*_*max*_ denotes current normalized to the max current elicited by 1000 µM Ca^2+^ at given pH_i_, [Ca^2+^] denotes free Ca^2+^ concentration, *H* denotes Hill coefficient, and *EC*_*50*_ denotes the half-maximal activation concentration of Ca^2+^.

Bell-shape fitting was performed with bell-shaped dose-response in Prism software (GraphPad). Briefly, the pH_i_ sensivivity (Y) and Ca^2+^ concentration (X) were fitted with the following equations,

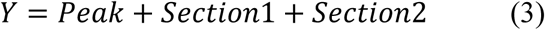

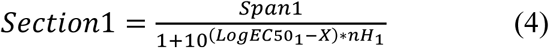

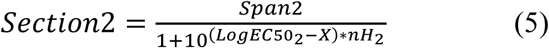

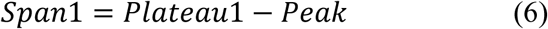

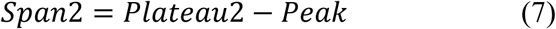

where, *Plateau1* and *Plateau 2* denote the plateaus at the left and right ends of the curve, which are set to be 0 in this study; *Peak* is the maximum value of the curve. *X* is the Ca^2+^ concentration. *LogEC50_1* and *LogEC50_2* are the concentrations that give half-maximal stimulatory and inhibitory effects in the same units as *X*; *nH*_*1*_ and *nH*_*2*_ are the unitless slope factors or Hill slopes, which are set to be 1 in this study.

PDB coordinate files were downloaded from the Protein Data Bank website https://www.rcsb.org/. All figures were generated in Pymol software (Schrödinger, Inc.).

### Quantifying phospholipid scrambling activity

To analyze the accumulation of AnV fluorescent signal on cell surfaces overtime, a previously reported MATLAB (Mathworks) code was used (T. Le *et al*., 2019). Briefly, a region of interest (ROI) was manually chosen around the scrambling cells, and the AnV fluorescent intensity was calculated using the following equation for each frame,

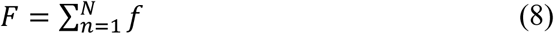

where *f* equals the fluorescent intensity of each pixel and *N i*s the number of the pixels in the ROI.

The time-dependent increase of AnV fluorescence was fitted with generalized logistic equation (Yin *et al*., 2003),

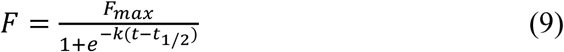

where F_max_ is the maximum normalized AnV fluorescent intensity, which is set to 100% in this study; k is the maximum growth rate or the ‘slope’ of the ‘linear phase’ in the sigmoidal curve; t_1/2_ is the time point at which the growth rate reaches its maximum and the fluorescent intensity reaches 1/2 of its maximum value.

### Statistical analysis

All statistical analyses were performed with Clampfit 10.7, Excel and Prism software (GraphPad). Two-tailed Student’s t-test was used for single comparisons between two groups (paired or unpaired), and one-way ANOVA following by Tukey’s test was used for multiple comparisons. Data were represented as mean ± standard error of the mean (SEM). Symbols *, **, ***, **** and ns denote statistical significance corresponding to p-value <0.05, <0.01, <0.001, <0.0001 and no significance, respectively.

### Data Availability Statement

All data discussed in the paper will be made available to readers upon request.

## Acknowledgement

We are grateful to Son C. Le for his help on electrophysiology and plotting the structural models. We thank Drs. Jianmin Cui and Jorg Grandl their constructive suggestions on the projects. We appreciate Trieu Le, Yang Zhang and Ping Dong for their generous help and comments on this project. This work was supported by NIH-DP2-GM126898 (to H.Y.).

## Author Contributions

H.Y. conceived and supervised the project. P.L. performed phosopholipid scrambling and electrophysiology experiments, as well as data analysis. P.L. and H.Y. wrote the manuscript.

## Supplementary information

**Figure S1.**
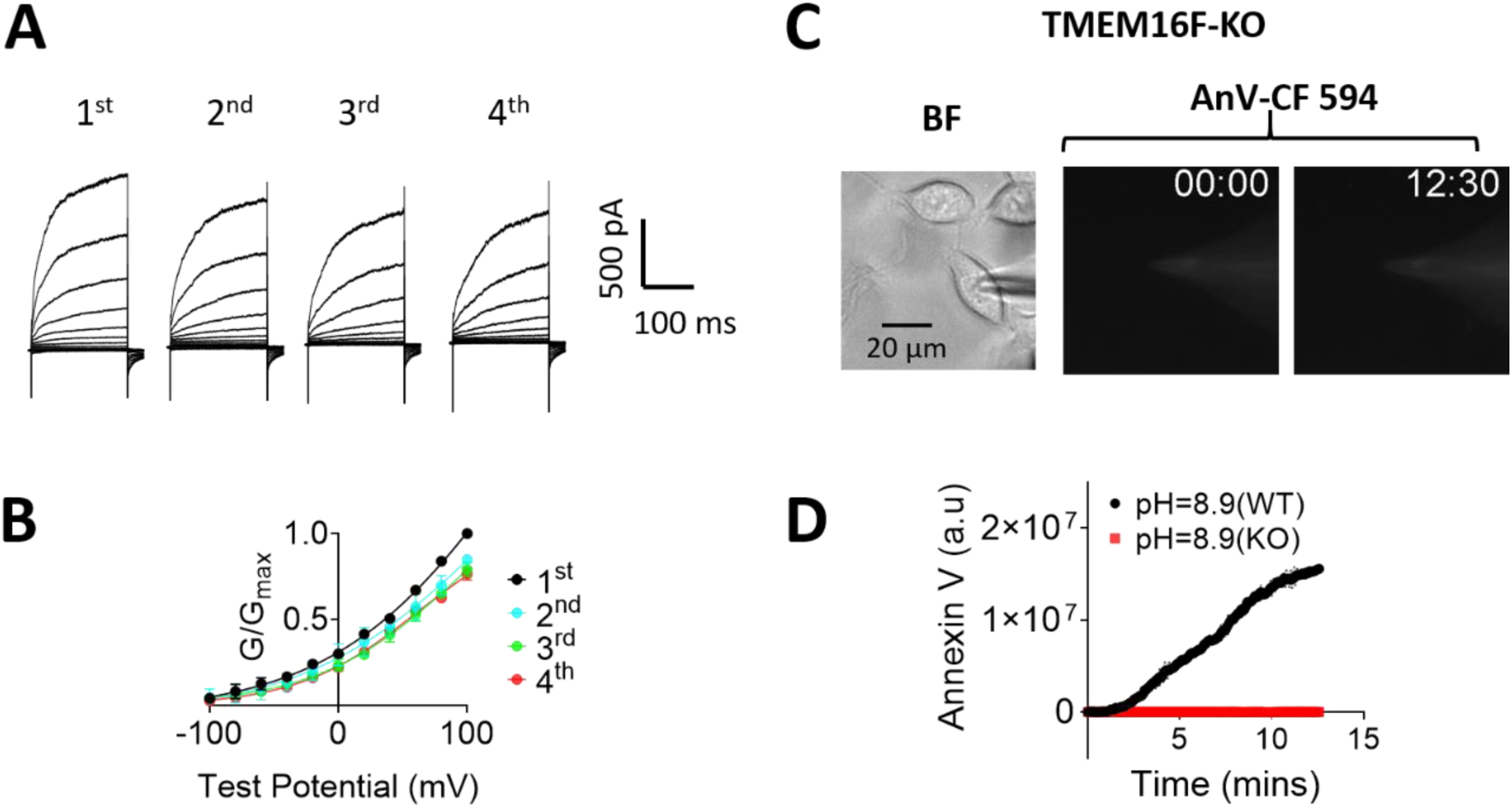
TMEM16F shows less rundown using voltage-step protocol and the knockout (KO) HEK293T cells show no scrambling activity even at high pH_i_. **(A)** Representative TMEM16F currents recorded from inside-out patches perfused with intracellular solutions containing 100 µM Ca^2+^ at pH_i_ 7.3. The interval between each trace was 20 seconds and all the recordings were from the same patch. **(B)** The *G-V* relationship of currents shown in **(A)**. Error bars represent mean ± SEM (n=3). **(C)** Representative images of lipid scrambling in TMEM16F-KO HEK293T cells. The patch pipette contained 100 μM free Ca^2+^ at pH_i_=8.9. No CaPLSase activity was observed in TMEM16F-KO HEK293T cells. **(D)** Representative fluorescent intensity of AnV binding for TMEM16F-eGFP stable HEK293T and TMEM16F-KO HEK293T cells at pH_i_=8.9 (n=4).

**Figure S2.**
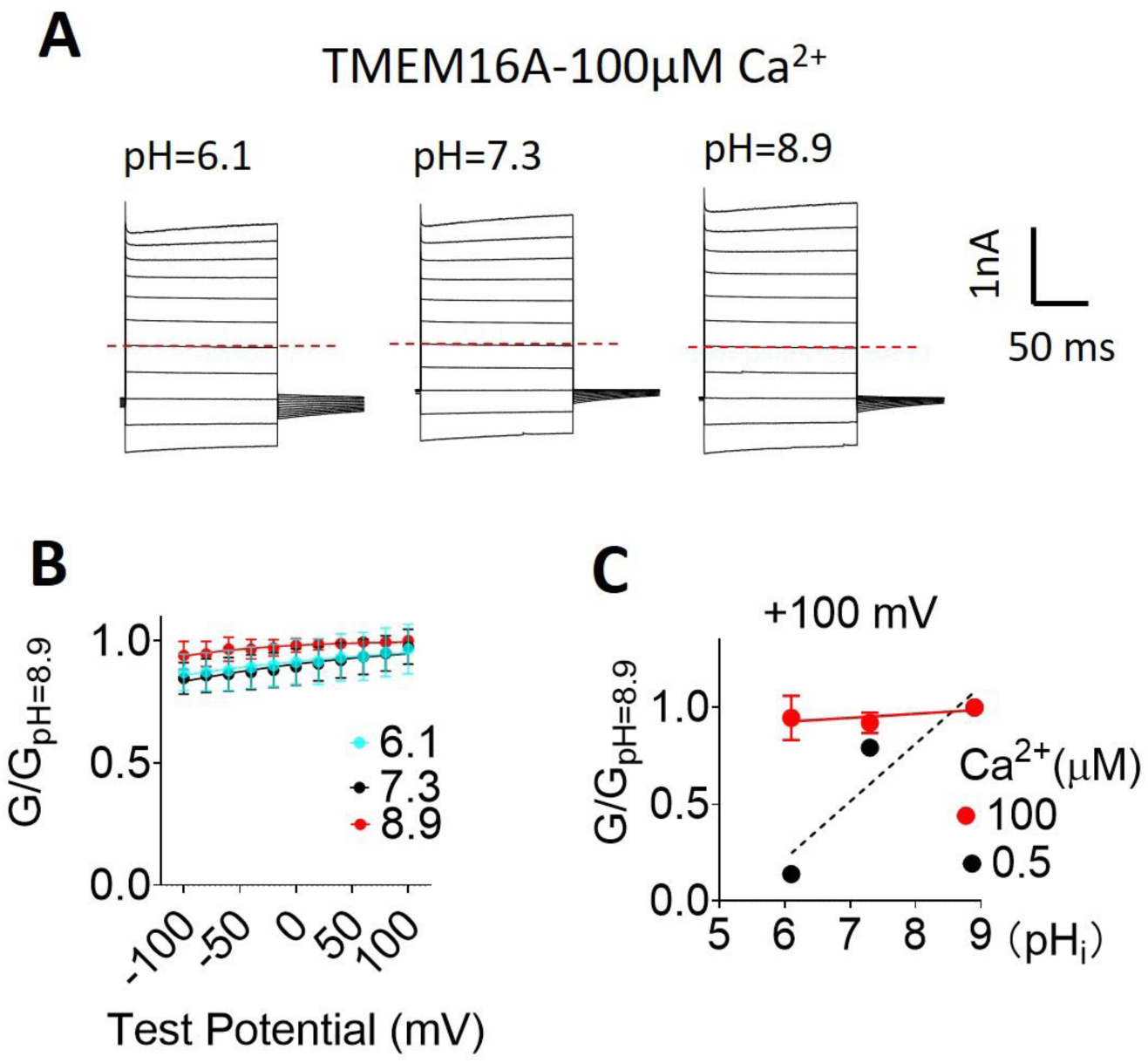
TMEM16A-CaCC losses pH_i_ regulation under saturating pH_i_. **(A)** Representative TMEM16A currents recorded from inside-out patches perfused with intracellular solutions containing 100 µM Ca^2+^ at different pH_i_. (B) *G-V* relationships at different pHi under 100 µM Ca^2+^. All conductance were normalized to the maximum conductance at pH_i_=8.9 and +100 mV. Error bar represents standard deviation (n=5). **(C)** *G-pH*_*i*_ curve of TMEM16A at 100 µM Ca^2+^ (red, solid line) and 0.5 µM Ca^2+^ (black, dotted line). The slopes from linear fits are 0.02 and 0.3, respectively. Error bar represents standard deviation (n=5).

**Figure S3.**
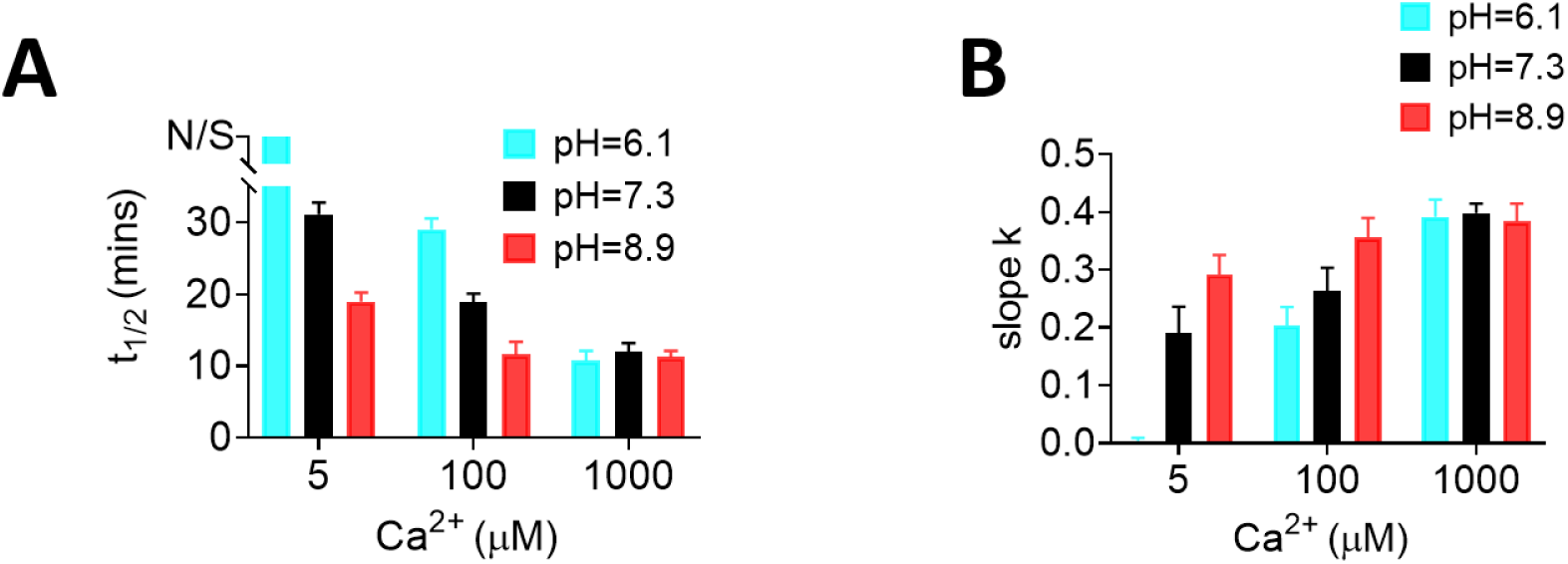
pH_i_ regulation on TMEM16F scrambling activity is Ca^2+^dependent. **(A)** The t_1/2_ of lipid scrambling at different pH_i_ under different Ca^2+^ concentrations including 5, 100 and 1000 μM. N/S represents no scrambling detected at given time. Error bars represent SEM (n =5). **(B)** The slopes from logistic equation fitting at different pH_i_ under different Ca^2+^ concentrations including 5, 100 and 1000 μM. Error bars represent SEM (n =5).

**Figure S4.**
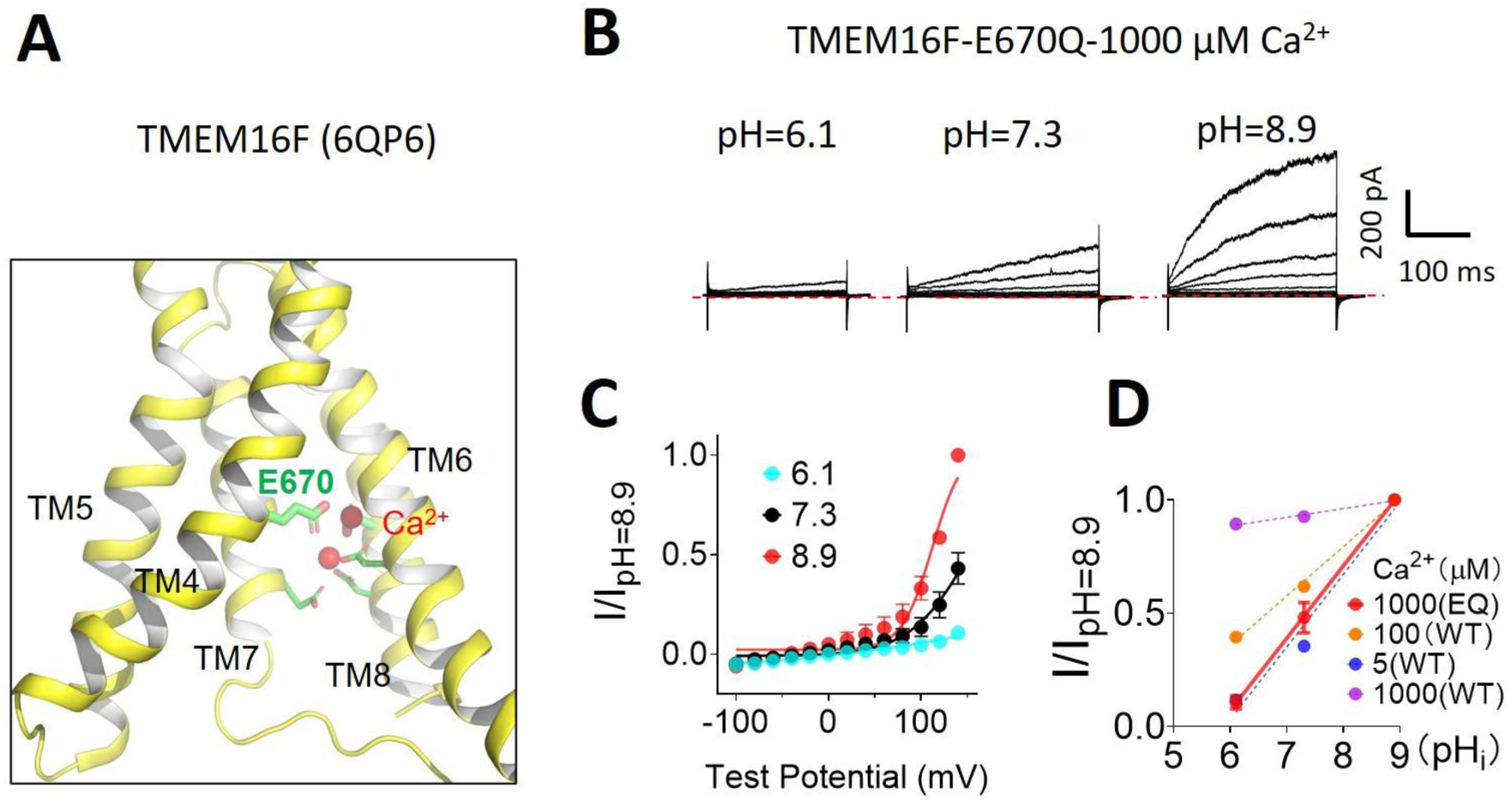
The Ca^2+^ binding site mutation E670Q enhances pH_i_ effects on TMEM16F channel activation. (**A**) Location of the Ca^2+^ binding residue E670. **(B**) Representative TMEM16F-E667Q currents recorded from inside-out patches perfused with intracellular solutions containing 1000 µM Ca^2+^ at different pHi. Currents were elicited by voltage steps from −100 to +140 mV with 20 mV increments. The holding potential was −60 mV. (**C**) *I-V* relationship of E670Q mutant channel recorded at different pHi. (**D**) *I-pHi* relationship of E670Q (EQ) under 1000 µM Ca^2+^ (red, solid line) with averaged slope of 0.32. *G-pH*_*i*_ relationships of WT TMEM16F at different Ca^2+^ concentrations were plotted as dashed line as references.

**Figure S5.**
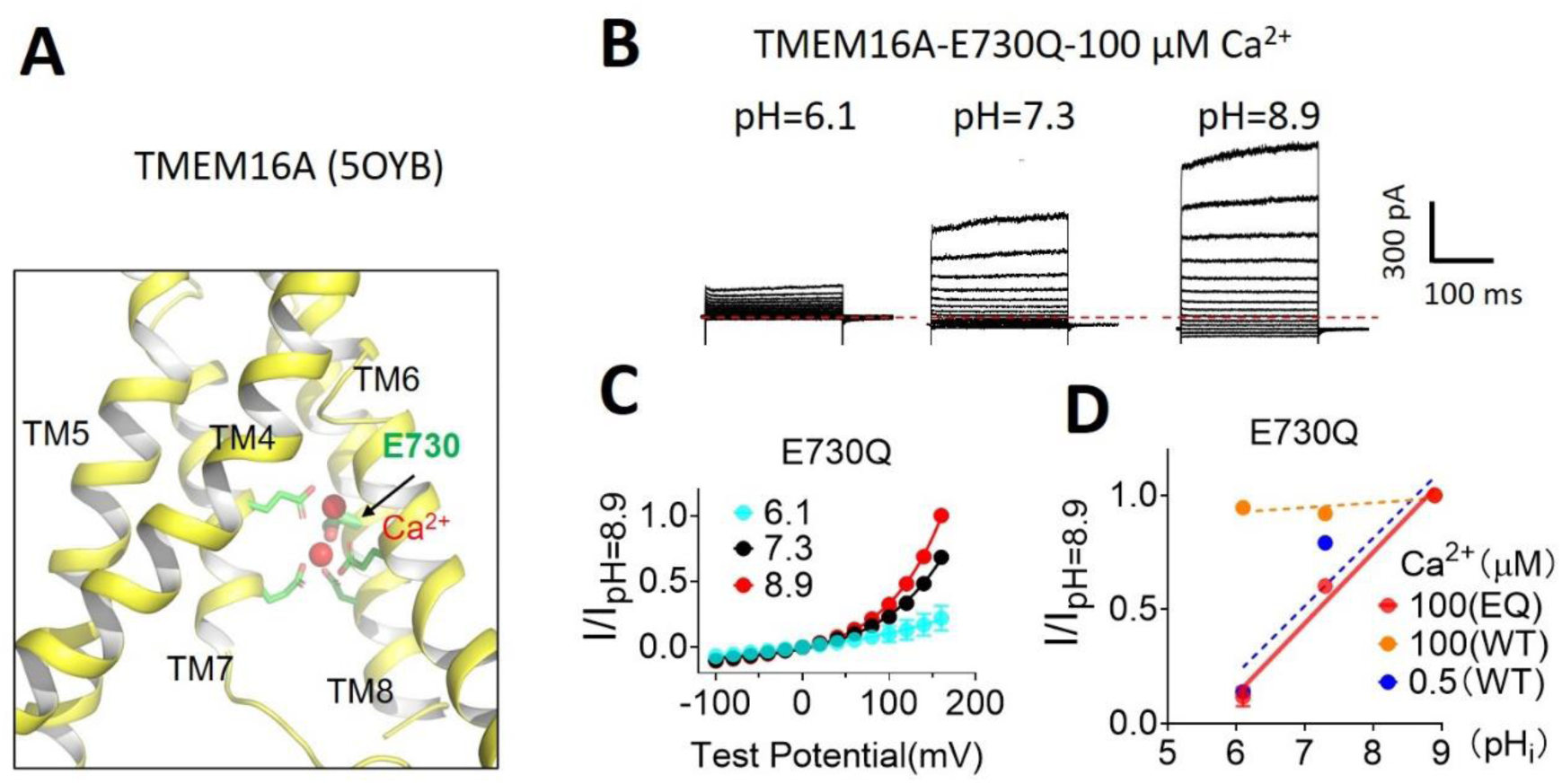
The Ca^2+^ binding site mutation E730Q enhances pH_i_ effects on TMEM16A activation. (**A**) Location of the Ca^2+^ binding residue E730 (PDB ID: 6BGI). **(B**) Representative TMEM16A-E730Q currents recorded from inside-out patches perfused with intracellular solutions containing 100 µM Ca^2+^ at different pH_i_. Currents were elicited by voltage steps from −100 to +160 mV with 20 mV increments. The holding potential was −60 mV. (**C**) *I-V* curve of E730Q current recorded at different pH_i_. (**D**) *I-pH*_*i*_ relationship of TMEM16A E730Q (EQ) under 100 µM Ca^2+^ (red, solid line) with averaged slope of 0.31. *G-pH*_*i*_ relationships of WT TMEM16A at different Ca^2+^ concentrations were plotted as dashed line as references.

**Figure S6.**
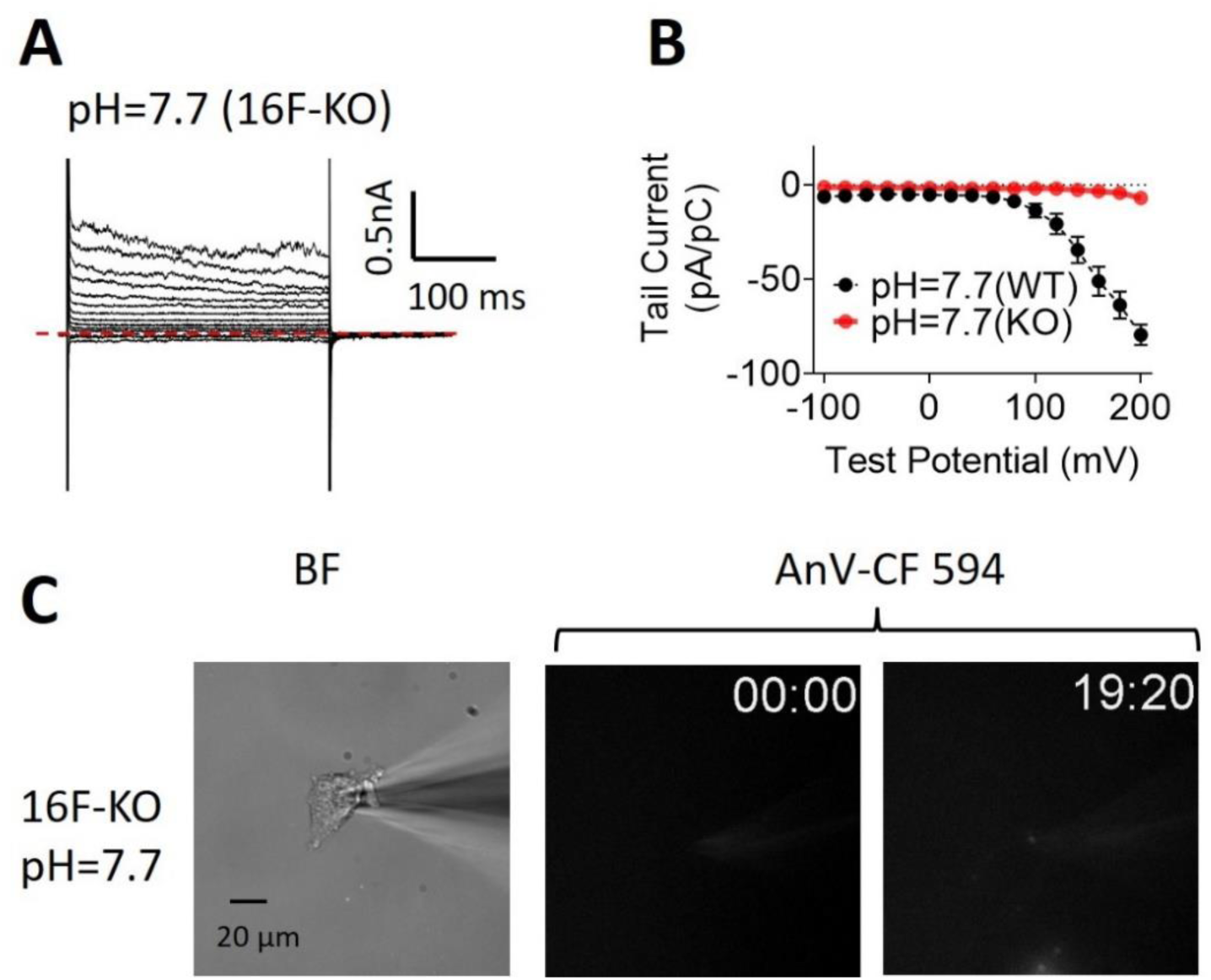
TMEM16F knockout (KO) BeWo cells lack TMEM16F-current and lipid scrambling. **(A)** Whole-cell current in TMEM16F-KO BeWo cells with pipette solution containing 5 µM Ca^2+^ at pH_i_=7.7. Currents were elicited by voltage steps from −100 to +200 mV with 20 mV increments. The holding potential was −60 mV. Compared with endogenous TMEM16F current in BeWo cells (Fig. 8A), the outward rectifying current of unknown identity in the TMEM16F KO BeWo cells is small in amplitude and lacks tail current. **(B)** The *I-V* relationship constructed using tail currents in (A). Error bar represents SEM (n=5 for WT and n=4 for KO). **(C)** Representative lipid scrambling activity of the TMEM16F-KO BeWo cells at different time points (right, top corner, minutes followed by seconds). None of the cells patched showed PS exposure (n=4).

### Captions for Movies S1-S6

**Movie S1**. Lipid Scrambling activity of HEK293T cells overexpressing TMEM16F with 100 µM Ca^2+^ at pH_i_=6.1, pH_i_=7.3 and pH_i_=8.9 (Related to Fig. 2B).

**Movie S2**. Lipid scrambling activity of TMEM16F-KO HEK293T cells with 100 µM Ca^2+^ at pH_i_=8.9 (Related to Fig. S1).

**Movie S3**. Lipid scrambling activity of HEK293T cells overexpressing TMEM16F with 5 µM Ca^2+^ at pH_i_=6.1, pH_i_=7.3 and pH_i_=8.9 (Related to Fig. 5A).

**Movie S4**. Lipid scrambling activity of HEK cells overexpressing TMEM16F with 1000 µM Ca^2+^ at pH_i_=6.1, pH_i_=7.3 and pH_i_=8.9 (Related to Fig. 5E).

**Movie S5**. Lipid scrambling activity of BeWo cells with 5 µM Ca^2+^ at pH_i_=7.2 and pH_i_=7.7 (Related to Fig.8D).

**Movie S6**. Lipid scrambling activity of TMEM16F knockout (KO) BeWo cells with 5 µM Ca^2+^ at pH_i_=7.7 (Related to Fig. S6).

